# Scalable microbial strain inference in metagenomic data using StrainFacts

**DOI:** 10.1101/2022.02.01.478746

**Authors:** Byron J. Smith, Xiangpeng Li, Zhou Jason Shi, Adam Abate, Katherine S. Pollard

## Abstract

While genome databases are nearing a complete catalog of species commonly inhabiting the human gut, their representation of intraspecific diversity is lacking for all but the most abundant and frequently studied taxa. Statistical deconvolution of allele frequencies from shotgun metagenomic data into strain genotypes and relative abundances is a promising approach, but existing methods are limited by computational scalability. Here we introduce StrainFacts, a method for strain deconvolution that enables inference across tens of thousands of metagenomes. We harness a “fuzzy” genotype approximation that makes the underlying graphical model fully differentiable, unlike existing methods. This allows parameter estimates to be optimized with gradient-based methods, speeding up model fitting by two orders of magnitude. A GPU implementation provides additional scalability. Extensive simulations show that StrainFacts can perform strain inference on thousands of metagenomes and has comparable accuracy to more computationally intensive tools. We further validate our strain inferences using single-cell genomic sequencing from a human stool sample. Applying StrainFacts to a collection of more than 10,000 publicly available human stool metagenomes, we quantify patterns of strain diversity, biogeography, and linkage-disequilibrium that agree with and expand on what is known based on existing reference genomes. StrainFacts paves the way for large-scale biogeography and population genetic studies of microbiomes using metagenomic data.

## Introduction

Intra-specific variation in microbial traits are widespread and are biologically important in human associated microbiomes. Strains of a species may differ in their pathogenicity (Loman et al., 2013), antibiotic resistance (Shoemaker et al., 2001), impacts on drug metabolism (Haiser et al., 2014), and ability to utilize dietary components (Patrick et al.; Martens et al., 2021). Standard methods for analysis of complex microbial communities are limited to coarser taxonomic resolution due to their reliance on slowly evolving marker genes (Case et al., 2007-January) or on genome reference databases lacking diverse strain representation (Nayfach et al., 2020). Approaches that quantify microbiomes at the level of strains may better capture variation in microbial function (Albanese and Donati, 2017), provide insight into ecological and evolutionary processes (Garud and Pollard, 2019), and discover previously unknown microbial etiologies for disease (Yan et al., 2020).

Shotgun metagenomic data can in principle be used to track strains by looking for distinct patterns of alleles observed across single nucleotide polymorphisms (SNPs) within the species. Several tools have recently been developed that count the number of metagenomic reads containing alleles across SNP sites (Nayfach et al., 2016; Costea et al., 2017b; Truong et al., 2017; Beghini et al., 2021; Olm et al., 2021; Shi et al., 2021). Comparisons of the resulting “metagenotypes” across samples has been used to track shared strains (Li et al., 2016; Olm et al., 2021), or to interrogate the biogeography (Costea et al., 2017a; Truong et al., 2017) and population genetics of species (Garud et al., 2019). The application of this approach is limited, however, by low sequencing coverage, which results in missing values at some SNP sites, and co-existing mixtures of strains, which introduce ambiguity about the taxonomic source of each metagenomic read.

One promising solution to these challenges is statistical strain deconvolution, which harnesses multiple metagenotypes (e.g., a collection of related samples) to simultaneously estimate the genotypes and relative abundances of strains across samples. Several tools have been developed that take this approach, including Lineage (O’Brien et al., 2014), Strain Finder (Smillie et al., 2018), DESMAN (Quince et al., 2017), and ConStrains (Luo et al., 2015). These methods have been used to track the transmission of inferred strains from donors’ to recipients’ microbiomes after fecal microbiota transplantation (FMT) (Smillie et al., 2018; Chu et al., 2021; Watson et al., 2021; Smith et al., 2022). The application of strain deconvolution has been limited, however, by the computational demands of existing methods, where runtimes scale poorly with increasing numbers of samples, latent strains, and SNPs considered. One reason for this poor scaling is the discreteness of alleles at each SNP, which has led existing methods to use expectation maximization algorithms to optimize model parameters (Smillie et al., 2018), or Markov chain Monte Carlo to sample from a posterior distribution (O’Brien et al., 2014; Luo et al., 2015; Quince et al., 2017).

Here we take a different approach, extending the strain deconvolution framework by relaxing the discreteness constraint and allowing genotypes to vary continuously between alleles. The use of this “fuzzy” genotype approximation makes our underlying model fully differentiable, and allows us to apply modern, gradient-based optimization algorithms to estimate strain genotypes and abundances. Here we show that the resulting tool, StrainFacts, can scale to tens of thousands of samples, hundreds of strains, and thousands of SNPs, opening the door to strain inference in large metagenome collections.

## Materials and Methods

### A fully differentiable probabilistic model of metagenotype data

#### Metagenotypes

A metagenotype is represented as a count matrix of the number of reads with each allele at a set of SNP sites for a single species in each sample. This can be gathered directly from metagenomic data, for instance by aligning reads to a reference genome and counting the number of reads with each allele at SNP sites. In this study we use GT-Pro (Shi et al., 2021), which instead counts exact k-mers associated with known single nucleotide variants. Although the set of variants at a SNP may include any of the four bases, here we constrain metagenotypes to be biallelic: reference or alternative. For a large majority of SNPs, only two alleles are observed across reference genomes (Shi et al., 2021). Metagenotypes from multiple samples are subsequently combined into a 3-dimensional array.

#### Deconvolution of metagenotype data

StrainFacts is based on a generative, graphical model of biallelic metagenotype data (summarized in Supplementary Fig. S1) which describes the allele frequencies at each SNP site in each sample (*p*_*ig*_ for sample *i* and SNP *g*) as the product of the relative abundance of strains 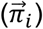 and their genotypes, *γ*_*sg*_, where 0 indicates the reference and 1 indicates the alternative allele for strain *s*. This functional relationship is therefore *p*_*ig*_ = ∑_*s*_ *γ*_*sg*_ × *π*_*is*_, In matrix form, equivalently, we notate this as **P = ΓΠ** (Table 1).

**Table 1:** Symbols used to describe the StrainFacts model

| Symbols | Description |
| --- | --- |
| $i = 1, \dots, N$ | Index and number of samples |
| $s = 1, \dots, S$ | Index and number of strains |
| $g = 1, \dots, G$ | Index and number of SNP sites |
| $y_{ig}, m_{ig}$ | Counts of reads with the alternative allele; the total count of both reference and alternative alleles at SNP $g$ in sample $i$ |
| $p_{ig}$ | Alternative allele frequency at SNP $g$ in sample $i$ |
| $\gamma_{sg}, \vec{\gamma}_g$ | Allele at SNP $g$ in strain $s$; vector of alleles for all strains |
| $\pi_{is}, \vec{\pi}_i$ | Relative abundance of strain $s$ in sample $i$; vector of relative abundances for all strains |
| $\epsilon_i$ | Sequencing error rate in sample $i$ |
| $\alpha$ | Concentration parameter of the BetaBinomial distribution |
| $\vec{\rho}$ | Metacommunity strain composition |
| $\mathbf{Y}, \mathbf{M}, \mathbf{P}, \mathbf{\Gamma}, \mathbf{\Pi}$ | Matrices composed of the above elements |

The crux of strain deconvolution is taking noisy observations of **P**—based on the observed alternative allele counts **Y** and total counts **M** obtained from metagenotypes across multiple samples—and determining suitable matrices *Γ* and **Π**. This notation highlights parallels to non-negative matrix factorization (NMF). Like NMF, given a choice of loss function, *L*, this inference task can be transformed into a constrained optimization problem, where 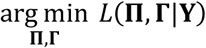 is a scientifically useful estimate of these two unobserved matrices. We take the approach of explicitly modeling the stochasticity of observed metagenotypes, placing priors on **Π** and **Γ**, and taking the resulting posterior probability as the loss function. This “maximum a posteriori” (MAP) approach has also been applied to NMF (Schmidt et al., 2009). However, unlike NMF, where the key constraint is that all matrices are non-negative, the metagenotype deconvolution model also constrains the elements of **P** and **Γ** to lie in the closed interval [0,1], and the rows of **Π** are are “on the *S* − 1-simplex”, i.e. they sum to one.

#### Fuzzy genotypes and the shifted-scaled Dirichlet distribution

StrainFacts does *not* constrain the elements of **Γ** to be discrete—i.e. in the set {0,1} for biallelic sites—in contrast to prior tools: DESMAN (Quince et al., 2017), Lineage (O’Brien et al., 2014), and Strain Finder’s (Smillie et al., 2018) exhaustive search. Instead, we allow genotypes to vary continuously in the open interval between fully reference (0) and fully alternative (1). The use of fuzzy-genotypes serves a key purpose: by replacing the only discrete parameter with a continuous approximation, our posterior function becomes fully differentiable, and therefore amenable to efficient, gradient-based optimization. When not using the exhaustive search strategy, Strain Finder also treats genotypes as continuous to accelerate inference, but these are discretized after each iteration. We show below that inference with StrainFacts is faster than with Strain Finder.

Since true genotypes are in fact discrete, we place a prior on the elements of **Γ** that pushes estimates towards zero or one and away from intermediate—ambiguous—values. Similarly, we put a hierarchical prior on **Π** that regularizes estimates towards lower strain heterogeneity within samples, as well as less strain diversity across samples. This makes strain inferences more parsimonious and interpretable. We harness the same family of probability distributions, the shifted-scaled Dirichlet distribution (SSD) (Monti et al., 2011), for all three goals. We briefly describe our rationale and parameterization of the SSD distribution in the Supplementary Methods.

For each element of **Γ** we set the prior 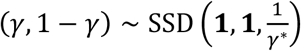. (Note that we trivially transform the 1-simplex valued (*γ*, 1 − *γ*) to the unit interval by dropping the second element.) Smaller values of the hyperparameter *γ*^*^ correspond to more sparsity in **Γ**. We put a hierarchical prior on **Π**, with the rows subject to the prior 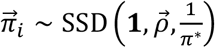 given a “metacommunity” hyperprior 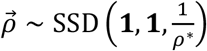, reflecting the abundance of strains across all samples. Decreasing the values of *γ*^*^, *ρ*^*^, and *π*^*^ increases the strength of regularization imposed by the respective priors.

#### Model specification

The underlying allele frequencies **P** are not directly observed due to sequencing error, and we include a measurement process in our model. We assume that the true allele is replaced with a random allele at a rate *ϵ*_*i*_ for all SNP sites *g* in sample *i*: 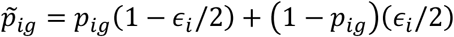. Given the total counts, **M**, we then model the observed alternative allele counts, **Y**,with the Beta-Binomial likelihood, parameterized with 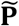 and one additional parameter—*α*^*^—controlling count overdispersion relative to the Binomial model.

To summarize, our model is as follows (in random variable notation; see Supplementary Fig. S1 for a plate diagram):

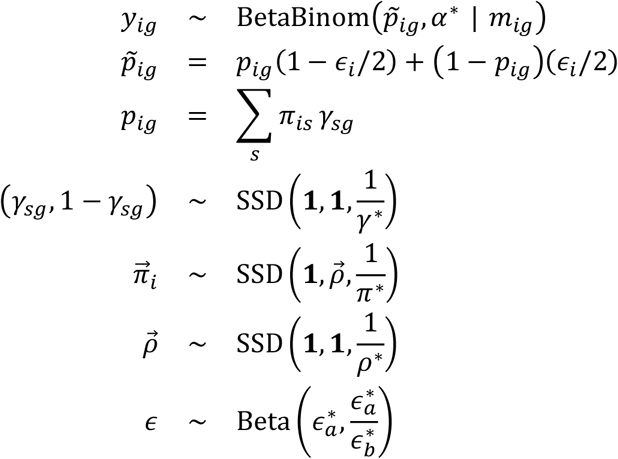

#### Model fitting

StrainFacts takes a MAP-based approach to inference on this model, using gradient-based methods to find parameter values that maximize the posterior probability of our model conditioned on the observed counts. We rely heavily on the probabilistic programming framework Pyro (Bingham et al., 2019), which is built on the PyTorch library (Paszke et al., 2019) for numerical methods.

Initial values for **Γ** and **Π** are selected using NMF, and all other parameters are initialized randomly (Supplementary Methods). In order to promote global convergence, we take a prior annealing approach (Supplementary Methods). While it is impossible to know in practice if we converge to a global optimum, we find that this procedure often leads to accurate estimates without the need for replicate fits from independent initializations.

#### Simulation and evaluation

Metagenotype data was simulated in order to enable direct performance benchmarking against ground-truth genotypes and strain compositions. For each independent simulation, discrete genotypes of length *G* for *S* strains were sampled as *S* × *G* independent draws from a symmetric Bernoulli distribution. The composition of strains in each of *N* samples were generated as independent draws from a Dirichlet distribution over *S* components having a symmetric concentration parameter of 0.4. Per-sample allele frequencies were generated as the product of the genotypes and the strain-composition matrices. Sequence error was set to *ϵ* = 0.01 for all samples. Finally metagenotypes at each SNP site were drawn from a Binomial 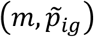 distribution, with a sequencing depth of *m* **=** 10 across all sites.

Estimates were evaluated against the simulated ground truth using five different measures of error (see Results).

#### Metagenotypes and reference genomes

We applied StrainFacts to data from two previously compiled human microbiome metagenomic datasets: stool samples from a fecal microbiota transplantation (FMT) study described in (Smith et al., 2022) and 20,550 metagenomes from a meta-analysis of publicly available data in (Shi et al., 2021). As described in that publication, metagenotypes for gut prokaryotic species were tallied using GT-Pro version 1.0.1 with the default database, which includes up to 1,000 of the highest quality genomes for each species from the Unified Human Gastrointestinal Genome (UHGG) V1.0 (Almeida et al., 2021). This includes both cultured isolates and high-quality metagenomic assemblies. This same database was used as a reference set to which we compared our inferred genotypes. Estimated genomic distances between SNPs were based on the UHGG representative genome.

We describe detailed results for *Escherichia coli_D* (id: 102506, MGYG-HGUT-02506), *Agathobacter rectalis* (id: 102492, MGYG-HGUT-02492), *Methanobrevibacter_A smithii* (id: 102163, MGYG-HGUT-02163), and CAG-279 sp1 (id: 102556, MGYG-HGUT-02556). These were selected to demonstrate application of StrainFacts to prevalent gram-positive and gram-negative bacteria in the human gut, the most prevalent archaeon, as well as an unnamed, uncultured, and largely unstudied species. We also describe detailed results for *Streptococcus thermophilus* (GT-Pro species id: 104345, representative UHGG genome: MGYG-HGUT-04345), selected for its high diversity in one sample of our single-cell sequencing validation.

#### Single-cell genome sequencing

Of the 159 samples with metagenomes described in the FMT study, we selected two samples for single-cell genomics (which we refer to as the “focal samples”). These samples were obtained from two different study subjects; one is a baseline sample and the other was collected after several weeks of FMT doses as described in (Smith et al., 2022). A full description of the single-cell genomics pipeline is included in the Supplementary Methods, and will be briefly summarized here. For each of the focal samples, microbial cells were isolated from whole feces by homogenization in phosphate buffered saline, 50 μm filter-based removal of large fecal particles, and density gradient separation. After isolating and thoroughly washing the density layer corresponding to the microbiota, this cell suspension was mixed with polyacrylamide precursor solution, and emulsified with a hydrofluoric oil. Aqueous droplets in oil were allowed to gellate before separating the resulting beads from the oil phase and washing. Beads were size selected to between 5 and 25 μm, with the goal of enriching for those encapsulated a single microbial cell. Cell lysis was carried out inside the hydrogel beads by incubating with zymolyase, lysostaphin, mutanolysin, and lysozyme. After lysis, proteins were digested with proteinase K, before thoroughly washing the beads. Tn5 tagmentation and barcode PCR were carried out using the MissionBio Tapestri microfluidics DNA workflow with minor modifications. After amplification, the emulsion was broken and the aqueous phase containing the barcoded amplicons was used for sequencing library preparation with Nextera primers including P5 and P7 sequences followed by Ampure XP bead purification. Libraries were sequenced by Novogene on an Illumina NovaSeq 6000.

Demultiplexed sequence data for each droplet barcode were independently processed with GT-Pro identically to metagenomic sequences. For each barcode, GT-Pro allele counts for a given species were assumed to be representative of a single strain of that species. These single-cell genotypes (SCGs) were filtered to those with >1% horizontal coverage over SNP sites, leaving 87 species with at least one SCG from either of the two focal samples. During analysis, a number of SCGs were found to have nearly identical patterns of horizontal coverage. These may have been formed by merging of droplets during barcoding PCR, which could have resulted in multiple barcodes in the same amplification. To reduce the impact of this artifact, allele counts from multiple SCGs were summed by complete-linkage, agglomerative clustering based on their depth profiles across SNP sites, at a 0.3 cosine dissimilarity threshold.

### Computational Analysis

#### Metagenotype filtering

From GT-Pro metagenotypes, we extracted allele counts for select species and removed SNPs that had <5% occurance of the minor allele across samples. Species with more than 5,000 SNPs after filtering, were randomly down-sampled without replacement to this number of sites. Samples with less than a minimum horizontal coverage (fraction of SNP sites with non-zero total counts), were also filtered out. This horizontal coverage threshold was set to 5% or 25% for the datasets from (Smith et al., 2022) or (Shi et al., 2021), respectively.

#### Strain Inference

For all analyses, StrainFacts was run with the following hyperparameters 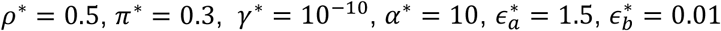. The learning rate was initially set to 0.05. Prior annealing was applied to both **Γ** and 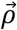 by setting *γ*^*^ and *ρ*^*^ to 1.0 and 5, respectively, for the first 2,000 steps of gradient descent, before exponentially relaxing these hyperparameters to their final values over the next 8,000 steps. After this annealing period, when parameters had not improved for 100 steps, the learning rate was halved until it had fallen below 10-6, at which point we considered parameters to have converged. These hyperparameters were selected through manual optimization and we found that they gave reasonable performance across the diverse datasets in this study.

The number of strains parameterized by our model was chosen as follows. For comparisons to SCGs, the number of strains was set at 30% of the number of samples—e.g. 33 strains were parameterized for *S. thermophilus* because metagenotypes from 109 samples remained after coverage filtering. For the analysis of thousands of samples described in (Shi et al., 2021), we parameterized our model with 200 strains and increased the numerical precision from 32 to 64 bits. After strain inference using the 5,000 subsampled SNPs, full-length genotypes were estimated post-hoc by conditioning on our estimate of **Π** and iteratively refitting subsets of all SNPs (Supplementary Methods).

For computational reproducibility we set fixed seeds for random number generators: 0 for all analyses where we only report one estimate, and 0, 1, 2, 3, and 4 for the five replicate estimates described for simulated datasets. Strain Finder was run with flags `--dtol 1 --ntol 2 -- max_reps 1`. We did not use `--exhaustive`, Strain Finder’s exhaustive genotype search strategy, as it is much more computationally intensive.

#### Genotype comparisons

Inferred fuzzy genotypes were discretized to zero or one for downstream analyses. SNP sites without coverage were treated as unobserved. Distances between genotypes were calculated as the masked, normalized Hamming distance, the fraction of alleles that do not agree, ignoring unobserved SNPs. Similarly, SCG genotypes and the metagenotype consensus were discretized to the majority allele. In comparing the distance between SCGs and these inferred genotypes sites missing from either the SCG or the metagenotype were treated as unobserved. Metagenotype entropy, a proxy for strain heterogeneity, was calculated for each sample as the depth weighted mean allele frequency entropy:

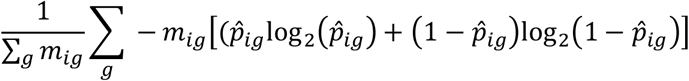

where 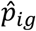 is the observed alternative allele frequency.

Where indicated, we dereplicated highly similar strains by applying average-neighbor agglomerative clustering at a 0.05 genotype distance threshold. Groups of these highly similar strains were replaced with a single composite strain with a genotype derived from the majority allele at each SNP site and assigned the sum of strain relative abundances in each sample. Subsequent co-clustering of these dereplicated inferred and reference strains was done in the same way, but at a 0.15 genotype distance threshold. After co-clustering, to test for enrichment of strains in “shared” clusters, we permuted cluster labels and re-tallied the total number of strains found in clusters with both inferred and reference strains. Likewise, to test for enrichment of “inferred-only” clusters we tallied the total number of strains found in clusters without reference strains after this shuffling. By repeating the permutation 9999 times, we arrived at an empirical null distribution to which we compared our true, observed values to calculate a P-value.

Pairwise linkage disequilibrium (LD) was calculated as the squared Pearson correlation coefficient across genotypes of dereplicated strains. Genome-wide 90th percentile LD, was calculated from a random sample of 20,000 or, if fewer, all available SNP positions. To calculate the 90th percentile LD profile, SNP pairs were binned at either an exact genomic distance or within a window of distances, as indicated. In order to encourage a smooth distance-LD relationship, windows at larger pairwise-distance spanned a larger range. Specifically the ith window covers the span [⌊10^(*i*−1)/*c*^⌋, ⌊10^*i*/*c*^⌋) where *c* **=** 30 so that 120 windows span the full range [1, 10^4^).

#### Software and code availability

StrainFacts is implemented in Python3 and is available at https://github.com/bsmith89/StrainFacts and v0.1 was used for all results reported here. Strain Finder was not originally designed to take a random seed argument, necessitating minor modifications to the code. Similarly, we made several modifications to the MixtureS (Li et al.) code allowing us to run it directly on simulated metagenotypes and compare the results to StrainFacts and Strain Finder outputs. Patch files describing each set of changes, as well as all other code and metadata needed to re-run our analyses are available at https://doi.org/10.5281/zenodo.5942586. For reproducibility, analyses were performed using Snakemake (Mölder et al., 2021) and with a Singularity container (Kurtzer et al., 2017) that can be obtained at https://hub.docker.com/repository/docker/bsmith89/compbio.

#### Runtime and memory benchmarking

Runtimes were determined using the Snakemake `benchmark:` directive, and memory requirements using the GNU `time` utility, version 1.8 with all benchmarks run on the Wynton compute cluster at the University of California, San Francisco. Across strain numbers and replicates, maximum memory usage for models with 10,000 samples and 1000 SNPs was, counterintuitively, less than for smaller models, likely because portions of runtime data were “swapped” to disk instead of staying in RAM. We therefore excluded data for these largest models from our statistical analysis of memory requirements.

## Results

### Scaling strain inference to hundreds of genotypes in thousands of samples

Inferring the genotypes and relative abundance of strains in large metagenome databases requires a deconvolution tool that can scale to metagenotypes with thousands of SNPs in tens-of-thousands of samples, while simultaneously tracking hundreds of microbial strains. To accomplish this we developed StrainFacts, harnessing fuzzy genotypes to accelerate inference on large datasets. We evaluated the practical scalability of the StrainFacts algorithm by applying it to simulated datasets of increasing size, and comparing its time and memory requirements to Strain Finder, a previously described method for strain inference. While several tools have been developed to perform strain deconvolution (e.g. Lineage O’Brien et al., 2014; and DESMAN Quince et al., 2017), Strain Finder’s model and approach to inference are the most similar to StrainFacts. We therefore selected it for comparison in order to directly assess the value of fuzzy genotypes.

We simulated five replicate metagenotypes for 120 underlying strains in 400 samples, and 250 SNPs, and then applied both StrainFacts and Strain Finder to these data parameterizing them with 120 strains. Both tools use random initializations, which can result in convergence to different optima. We therefore benchmarked runtimes for five independent initializations on each dataset, resulting in 25 total runs for each tool. In this setting, the median runtime for StrainFacts was just 17.2 minutes, while Strain Finder required a median of 6.4 hours. When run on a GPU instead of CPU, StrainFacts was able to fit these data in a median of just 5.1 minutes.

Since the correct strain number is not known a priori in real-world applications, existing strain inference tools need to be parameterized across a range of plausible strain counts, a step that significantly impacts runtime. To assess performance in this setting, we also fit versions of each model with 50% more strains than the ground-truth, here referred to as the “1.5x parameterization” in contrast to the 1x parameterization already described. In this setting, StrainFacts’ performance advantage was even more pronounced, running in a median of 17.1 minutes and just 5.3 minutes on GPU, while Strain Finder required 30.8 hours. Given the speed of StrainFacts, we were able to fit an even larger simulation with 2,500 samples and 500 strains. On a GPU, this took a median of 12.6 minutes with the 1x parameterization and, surprisingly, just 8.9 minutes with the 1.5x parameterization. We did not attempt to run Strain Finder on this dataset.

We next examined runtime scaling across a range of sample counts between 50 and 2,500. We applied Strain Finder and StrainFacts (both CPU and GPU) to simulated metagenotypes with 250 SNPs, and a fixed 1:5 ratio of strains to samples. Median runtimes for each tool at both the 1x and 1.5x parameterization demonstrate a substantially slower increase for StrainFacts as model size increases (Fig. 1A). Strain Finder was faster than StrainFacts on the 1x parameterization of a small simulation with 50 samples and 10 strains: 1.3 minutes median runtime versus 4 minutes for StrainFacts on a CPU and 2.8 minutes on a GPU. However, StrainFacts had faster median runtimes on all other datasets.

**Figure 1:**
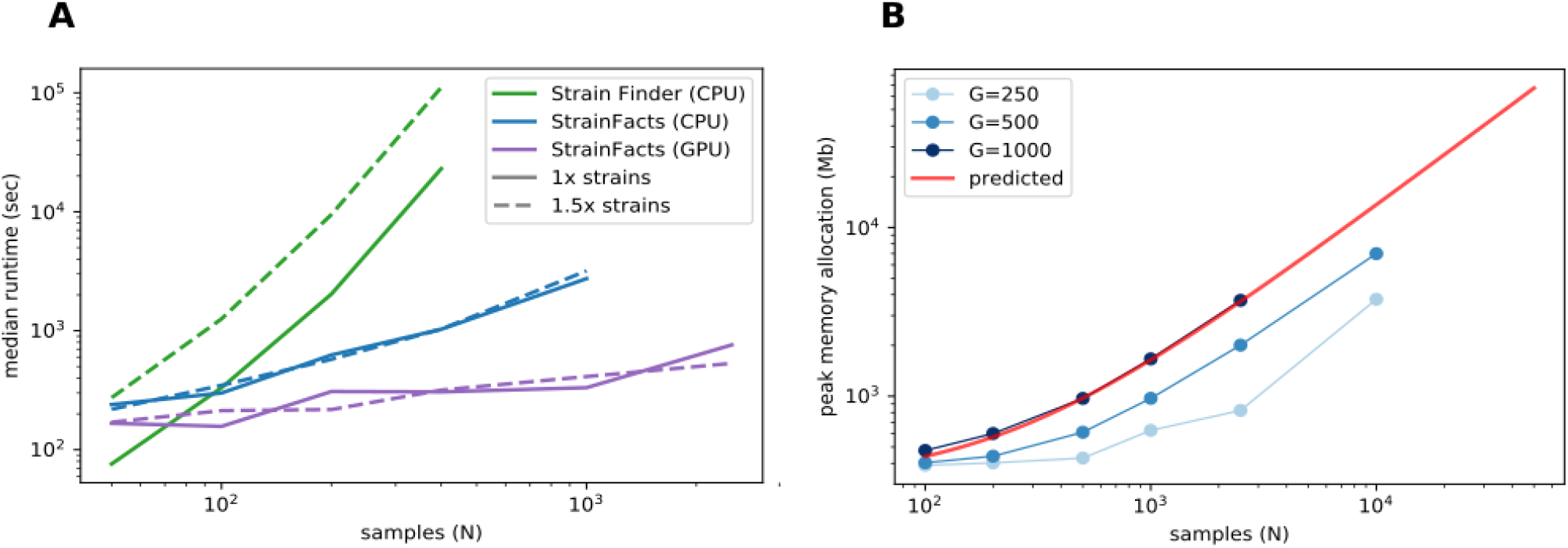
Computational scalability of strain inference on simulated data. **(A)** Runtime (in seconds, log scale) is plotted at a range of sample counts for both Strain Finder and StrainFacts, as well for the latter with GPU acceleration. Throughout, 250 SNPs are considered, and simulated strains are fixed at a 1:5 ratio with samples. Models are specified with this same number of strains (“1x strains”, solid lines) or 50% more (“1.5x strains”, dashed lines). Median of 25 simulation runs is shown. **(B)** Maximum memory allocation in a model with 100 strains is plotted for StrainFacts models across a range of sample counts (N) and SNP counts (G, line shade). Median of 9 replicate runs is shown. Maximum memory requirements are extrapolated to higher numbers of samples for a model with 1000 SNPs (red line). A version of this panel that includes a range of strain counts is included as Supplementary Fig. S2.

Given the good runtime scaling properties of StrainFacts, we next asked if computer memory constraints would limit its applicability to the largest datasets (Fig. 1A). A model fitting 10,000 samples, 400 strains, and 500 SNPs had a maximum memory allocation of 7.7 GB, indicating that StrainFacts’ memory requirements are satisfied on most contemporary CPU or GPU hardware and opening the door to even larger models. Using ordinary least squares, we fit the observed memory requirements to the theoretical, asymptomatic expectations, *𝒪*(*NS* + *NG* + *SG*), resulting in a regression R^2^ of 0.997. We then used this empirical relationship to extrapolate for even larger models (Fig. 1B), estimating that for a model of 400 strains and 1000 SNPs, 32 GB of memory would be able to simultaneously perform strain inference for more than 22,000 samples. This means StrainFacts can realistically analyse tens of thousands of samples on commercial GPUs.

### StrainFacts accurately reconstructs genotypes and population structure

We next set out to evaluate the accuracy of StrainFacts and to compare it to Strain Finder. We simulated 250 SNPs for 40 strains, generating metagenotypes across 200 samples. For both tools, we specified a model with the true number of strains, fit the model to this data, and compared inferences to the simulated ground-truth. For each of five replicate simulations we performed inference with five independent initializations, thereby gathering 25 inferences for each tool. As in (Smillie et al., 2018), we use the weighted UniFrac distance (Lozupone et al., 2007) as an integrated summary of both genotype and relative abundance error. By this index, StrainFacts and Strain Finder performed similarly well when applied to the simulated data (Fig. 2A). We repeated this analysis with the 1.5x parameterization to assess the robustness of inferences to model misspecification, finding that both tools maintained similar performance to the 1x parameterization. By comparison, considering too few strains (the 0.8x parameterization, fitting 32 strains) degraded performance dramatically for both tools, with StrainFacts performing slightly better. Thus, we conclude based on UniFrac distance that StrainFacts is as accurate as Strain Finder and that both models are robust to specifying too many strains.

**Figure 2:**
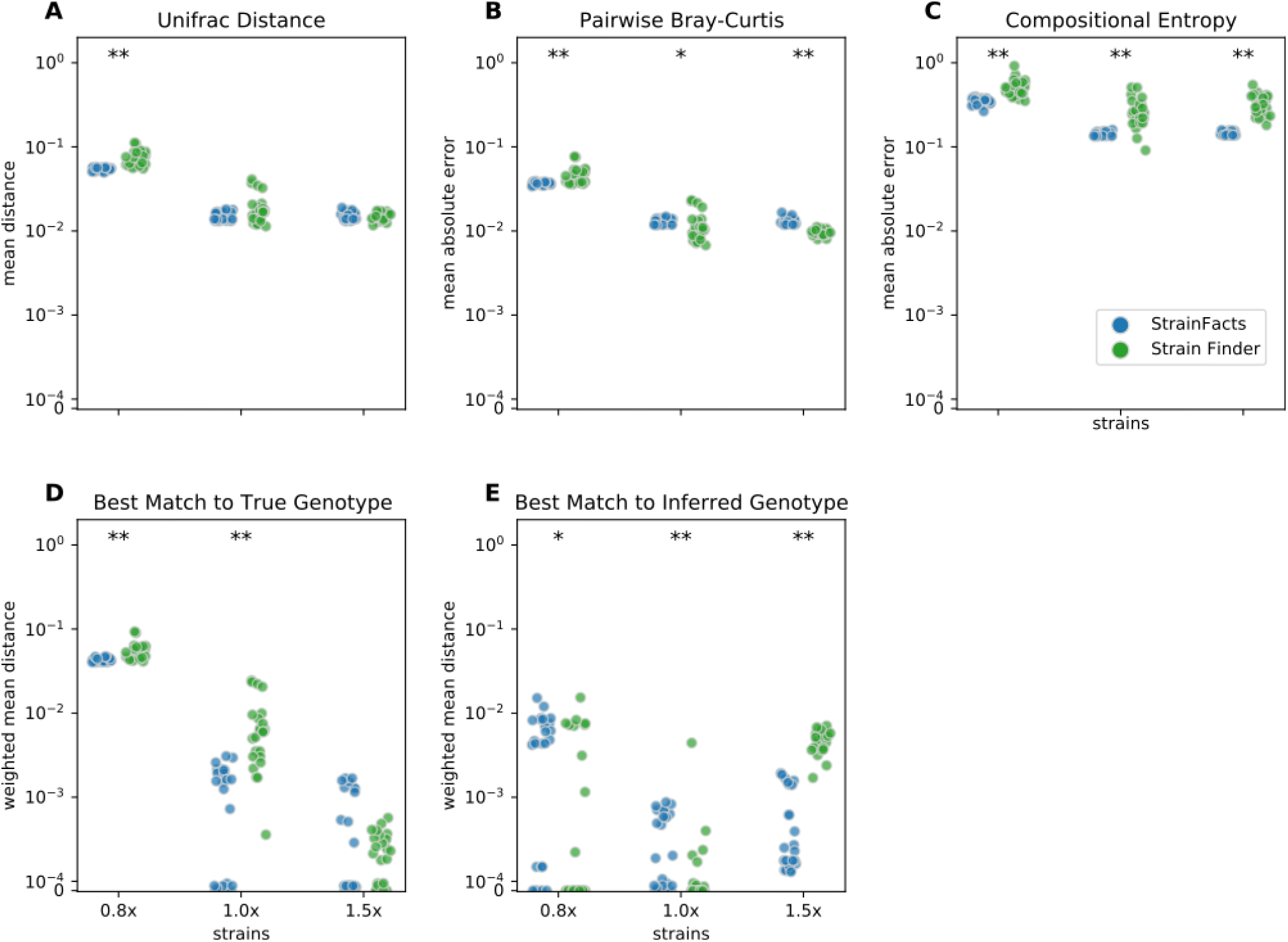
Accuracy of strain inference on simulated data. Performance of StrainFacts and Strain Finder are compared across five distinct accuracy indices, with lower scores reflecting better performance on each index. Simulated data had 200 samples, 40 underlying strains, and 250 SNPs. For each tool, 32, 40 and 60 strain models were parameterized (“0.8x”, “1x” and “1.5x” respectively), and every model was fit with five independent initializations to each simulation. All 25 estimates for each tool-parameterization combination are shown. Scores reflect **(A)** mean Unifrac distance between simulated and inferred strain compositions, **(B)** mean absolute difference between all-by-all pairwise Bray-Curtis dissimilarities calculated on simulated versus inferred strain compositions, **(C)** mean absolute difference in Shannon entropy calculated on simulated versus inferred strain compositions, **(D)** abundance weighted mean Hamming distance from each ground-truth strain to its best-match inferred genotype, and **(E)** the reverse: abundance weighted mean Hamming distance from each inferred strain to its best-match true genotype. Markers at the top of each panel indicate a statistical difference between tools at a p<0.05 (*) or p<0.001 (**) significance threshold by Wilcoxon signed-rank test. A version of this figure that includes accuracy comparisons to MixtureS is included as Supplementary Fig. S3.

To further probe accuracy, we quantified the performance of StrainFacts and Strain Finder with several other measures. First, we evaluated pairwise comparisons of strain composition by calculating the mean absolute error of pairwise Bray-Curtis dissimilarities (Fig. 2B). While, with the 1x parameterization, Strain Finder slightly outperformed StrainFacts on this index, the magnitude of the difference was small. This suggests that StrainFacts can be used for applications in microbial ecology that rely on measurements of beta-diversity.

Ideally, inferences should conform to Occam’s razor, estimating “as few strains as possible, but no fewer”. Unfortunately, Bray-Curtis error is not sensitive to the splitting or merging of co-abundant strains and UniFrac error is not sensitive to the splitting or merging of strains with very similar genotypes. To overcome this limitation, we calculated the mean absolute error of the Shannon entropy of the inferred strain composition for each sample (Fig. 2C). This score quantifies how accurately inferences reflect within-sample strain heterogeneity. StrainFacts performed substantially better on this score than Strain Finder for all three parameterizations, indicating more accurate estimation of strain heterogeneity.

Finally, we assessed the quality of genotypes reconstructed by StrainFacts compared to Strain Finder using the abundance weighted mean Hamming distance. For each ground-truth genotype, normalized Hamming distance is computed based on the best matching inferred genotype (Fig. 2D), then summarized as the mean weighted by the true strain abundance across all samples. We assessed the reverse as well: the abundance weighted mean, best-match Hamming distance for each inferred genotype among the ground-truth genotypes (Fig. 2E). These two scores can be interpreted as answers to the distinct questions “how well were the true genotypes recovered?” and “how well do the inferred genotypes reflect the truth?”, respectively. While StrainFacts and Strain Finder performed similarly on these indexes—which tool had higher accuracy varied by score and parameterization—StrainFacts’ accuracy was more stable across the 1x and 1.5x parameterizations. It should be noted that since strain genotypes are only inferred for SNP sites, the genome-wide genotype reconstruction error (which includes invariant sites) will likely be much lower than this Hamming distance. We examine the relationship between genotype distances and average nucleotide identity (ANI) in Supplementary Fig. S4 in order to contextualize our simulation results for those more familiar with ANI comparisons.

To expand our performance comparison to a second tool designed for strain inference, we also ran MixtureS on a subset of the simulations. MixtureS estimates strain genotype and relative abundance on each metagenotype individually and therefore does not leverage variation in strain abundance across samples. We found that it performed worse than Strain Finder and Strain Facts on the benchmarks (see Supplementary Fig. S3).

Overall, these results suggest that StrainFacts is capable of state-of-the-art performance with respect to several different scientific objectives in a realistic set of simulations. Performance was surprisingly robust to model misspecification with more strains than the simulation. Eliminating the computational demands of a separate model selection step further improves the scaling properties of StrainFacts.

### Single-cell sequencing validates inferred strain genotypes

Beyond simulations, we sought to confirm the accuracy of strain inferences in a real biological dataset subject to forms of noise and bias not reflected in the generative model we used for simulations. To accomplish this, we applied a recently developed, single-cell, genomic sequencing workflow to obtain ground-truth, strain genotypes from two fecal samples collected in a previously described, clinical FMT experiment (Smith et al., 2022) from two independent subjects. We ran StrainFacts on metagenotypes derived from these two focal samples as well as the other 157 samples in the same study.

Genotypes that StrainFacts inferred to be present in each of these metagenomes matched the observed SCGs, with a mean, best-match normalized Hamming distance of 0.039. Furthermore, the median distance was just 0.013, reflecting the outsized influence of a small number of SCGs with more extreme deviations. For many species, SCGs also match a consensus genotype—the majority allele at each SNP site in each metagenotype (see Fig. 3A). We found a mean distance to the consensus of 0.037 and a median of 0.009. Because this distance excludes sites without observed counts in the metagenotype, we masked these same sites in our inferred genotypes to uniformly contrast the consensus approach to StrainFacts genotypes. Overall, inferred genotypes were similar to the consensus, with a mean, masked distance of 0.031 (median of 0.009). However, the consensus approach fails for species with a mixture of multiple, co-existing strains. When we select only species with a metagenotype entropy of greater than 0.05, an indication of strain heterogeneity, we see that StrainFacts inferences have a distinct advantage, with a mean distance of 0.055 versus 0.069 for the consensus approach (median of 0.018 versus 0.022, p<0.001). These results validate inferred genotypes in a stool microbiome using single-cell genomics and demonstrate that StrainFacts accounts for strain-mixtures better than consensus genotypes do.

**Figure 3:**
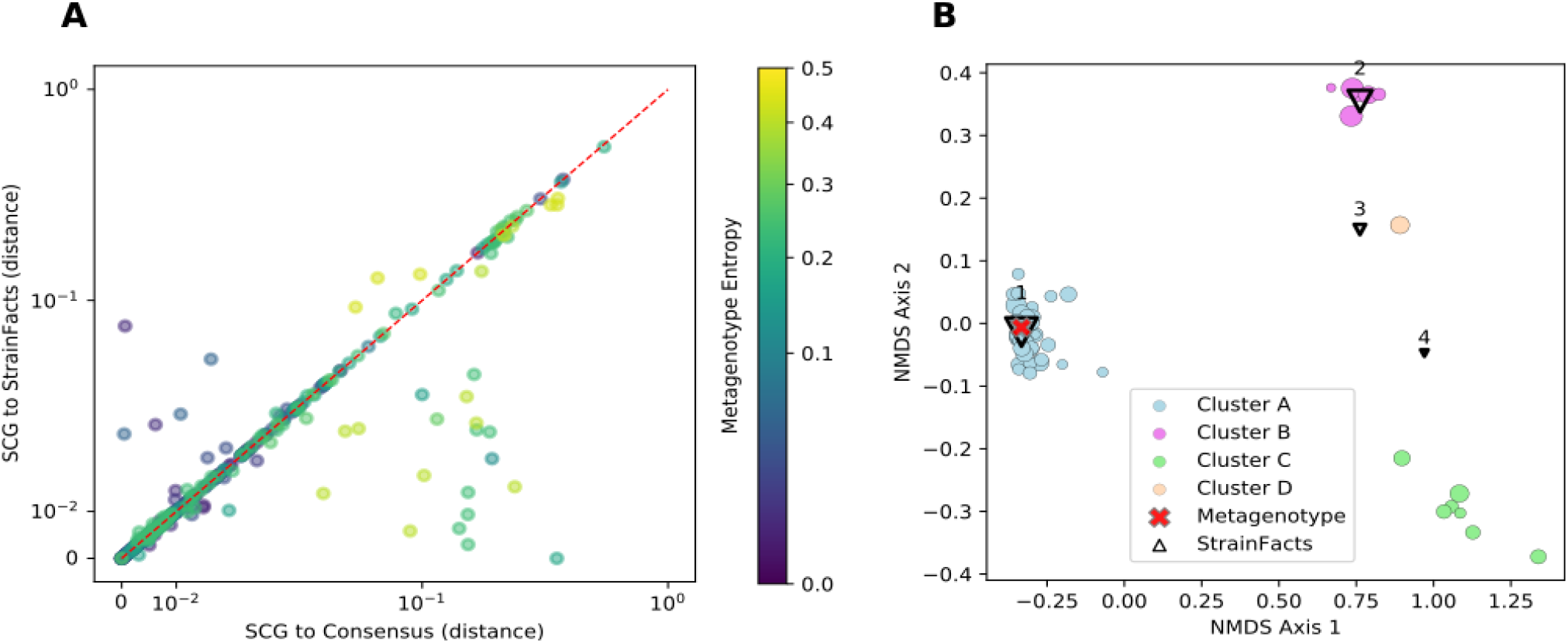
Inferred strains reflect genotypes from a single-cell sequencing experiment. **(A)** Distance between observed SCGs and StrainFacts inferences (X-axis) versus consensus genotypes (Y-axis). Points below and to the right of the red dotted line reflecting an improvement of our method over the consensus, based on the normalized, best-match Hamming distance. Each dot represents an individual SCG reflecting a putative genotype found in the analysed samples. SCGs from all species found in either of the focal samples are represented, and marker colors reflect the metagenotype entropy of that species in the relevant focal sample, a proxy for the potential strain diversity represented. Axes are on a “symmetric” log scale, with linear placement of values below 10^−2^. **(B)** A non-metric multidimensional scaling ordination of 68 SCGs and inferred genotypes for one species, S. thermophilus, with notably high strain diversity in one of the two focal samples. Circles represent SCGs, are colored by their assignment to one of four identified clusters, and larger markers indicate greater horizontal coverage. Triangles represent StrainFacts genotypes inferred to be at greater than 1% relative abundance, and larger markers reflect a higher inferred relative abundance. The red cross represents the consensus metagenotype of the focal sample.

Of the 75 species represented in our SCG dataset, one stood out for having numerous SCGs while reflecting a remarkably high degree of strain heterogeneity. Among 68 high-quality SCGs for *S. thermophilus*, cluster analysis identified four distinct types (here referred to as Clusters A - D), accounting for 48, 7, 6, and 1 SCGs, respectively (Fig. 3B). Independently, StrainFacts inferred four strains in the metagenomic data from the same stool sample, (Strain 1 - 4) with 57%, 32%, and 7%, and 3% relative abundance, respectively. We explored the concordance between clusters and StrainFacts inferences by assigning a best-match Hamming distance genotype among the inferred strains to each SCG (Table 2). For SCGs in three of the four clusters there was a low median distance to StrainFacts genotypes as well as a perfect 1-to-1 correspondence between strains and clusters. While this genotype concordance was broken for SCGs in cluster B, strain 4 was also inferred to be at the lowest relative abundance, suggesting that there may have been too little information encoded in the metagenotype data to accurately reconstruct that strain’s genotype. While SCG counts and inferred strain fractions do not match perfectly in this sample, this may be due to large differences between SCG and metagenomic sequencing technologies that could result in differentially biased sampling of strains. The SCG cluster with the largest membership was, however, matched with the strain inferred to be at the highest relative abundance. Our findings for *S. thermophilus* show that StrainFacts’ estimates of genotypes and relative abundances are remarkably accurate for samples with high strain heterogeneity, despite the challenges presented by real biological samples and low abundance strains.

**Table 2:**
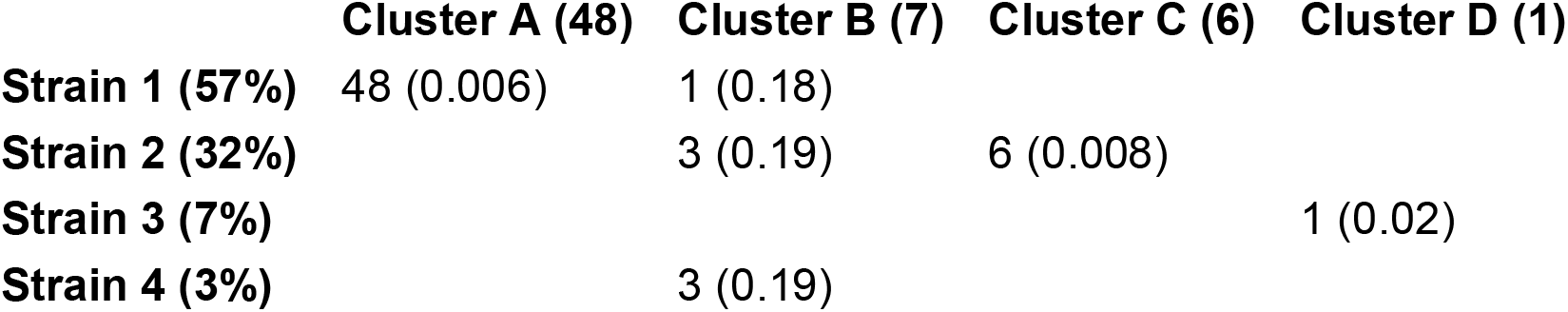
Concordance among SCGs of cluster assignments and the closest-match StrainFacts inferred genotype, among the four strains inferred to be at greater than 1% relative abundance in the analysed sample. The total number of SCGs in each cluster and the relative abundance of each inferred strain are indicated in parentheses in the column and row labels, respectively. Numbers in each cell indicate the number of SCGs at that intersection and values in parentheses indicate the median normalized Hamming distance of those SCGs to the inferred strain genotype.

### Analysis of genomic diversity using de novo strain inferences on thousands of samples

Having established the accuracy and scalability of StrainFacts, we applied it to a corpus of metagenotype data derived from 20,550 metagenomes across 44 studies, covering a large fraction of all publicly available human-associated microbial metagenomes (Shi et al., 2021). We performed strain inference on GT-Pro metagenotypes for four species: *Escherichia coli, Agathobacter rectalis, Methanobrevibacter smithii*, and CAG-279 sp1. *E. coli* and *A. rectalis* are two highly prevalent and well studied bacterial inhabitants of the human gut microbiome, and *M. smithii* is the most prevalent and abundant archaeon detected in the human gut (Scanlan et al., 2008). CAG-279, on the other hand, is an unnamed and little-studied genus and a member of the family *Muribaculaceae*. This family is common in mice (Lagkouvardos et al., 2019), but to our knowledge does not have representatives cultured from human samples.

For each species, we compared strains inferred by StrainFacts to those represented in the GT-Pro reference database, which is derived from the UHGG (Almeida et al., 2021). In order to standardize comparisons, we dereplicated inferred and reference strains at a 0.05 genotype distance threshold. Interestingly, dereplication had a negligible effect on StrainFacts results, reducing the number of *E. coli* strains by just 4 (to 119) with no reduction for the three other species. This suggests that the diversity regularization built into the StrainFacts model is sufficient to collapse closely related strains as part of inference.

As GT-Pro only tallies alleles at a fixed subset of SNPs, the relationship between genotype distances and ANI is not fixed. In order to anchor our results to this widely-used measure of genome similarity, we compared the genotype distance to genome-wide ANI for all pairs of genomes in the GT-Pro reference database for all four species. We find that the fraction of positions differing genome wide (calculated as 1 - ANI) was nearly proportional to the fraction of genotyped positions differing, but with a different constant of proportionality for each species: *E. coli* (0.0776, uncentered R^2^=0.994), *A. rectalis* (0.1069, R^2^=0.990), *M. smithii* (0.0393, R^2^=0.967), and CAG-279 (0.0595, R^2^=0.991). Additional details of this analysis can be found in Supplementary Fig. S4.

### StrainFacts recapitulates known diversity in well studied species

*E. coli, A. rectalis*, and *M. smithii* all have many genome sequences in GT-Pro reference database, presenting an opportunity to contrast inferred against reference strains. In order to evaluate the concordance between the two (Table 3 and Fig. 4), we co-clustered all dereplicated strains (both reference and inferred) at a 0.15 normalized Hamming distance threshold—note, crucially, that this distance reflects a much smaller full-genome dissimilarity, as it is based only on genome positions with polymorphism across metagenomes, ignoring conserved positions.

**Table 3:**
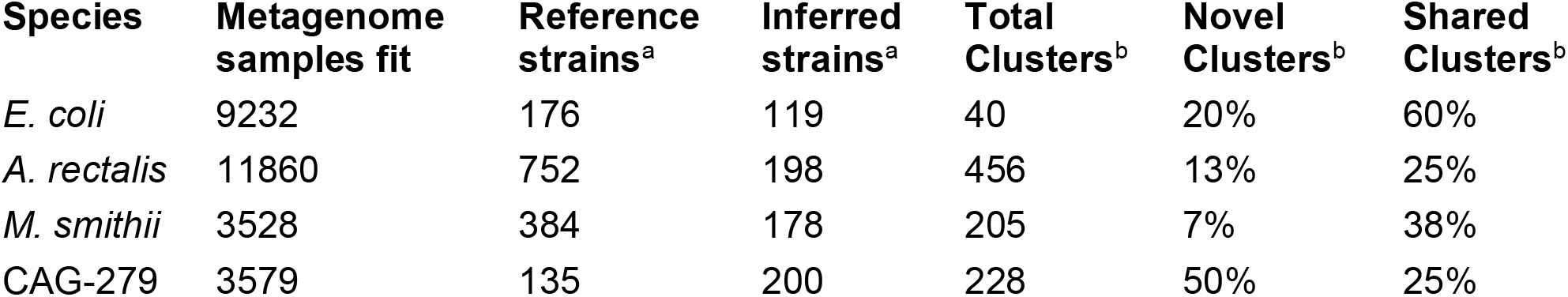
Dereplication and co-clustering of strains inferred from metagenomes or from a reference database ^a^ Dereplicated at 0.05 distance threshold ^b^ Co-clustered at a 0.15 distance threshold

| Species | Metagenome samples fit | Reference strains<sup>a</sup> | Inferred strains<sup>a</sup> | Total Clusters<sup>b</sup> | Novel Clusters<sup>b</sup> | Shared Clusters<sup>b</sup> |
| --- | --- | --- | --- | --- | --- | --- |
| *E. coli* | 9232 | 176 | 119 | 40 | 20% | 60% |
| *A. rectalis* | 11860 | 752 | 198 | 456 | 13% | 25% |
| *M. smithii* | 3528 | 384 | 178 | 205 | 7% | 38% |
| CAG-279 | 3579 | 135 | 200 | 228 | 50% | 25% |

**Figure 4:**
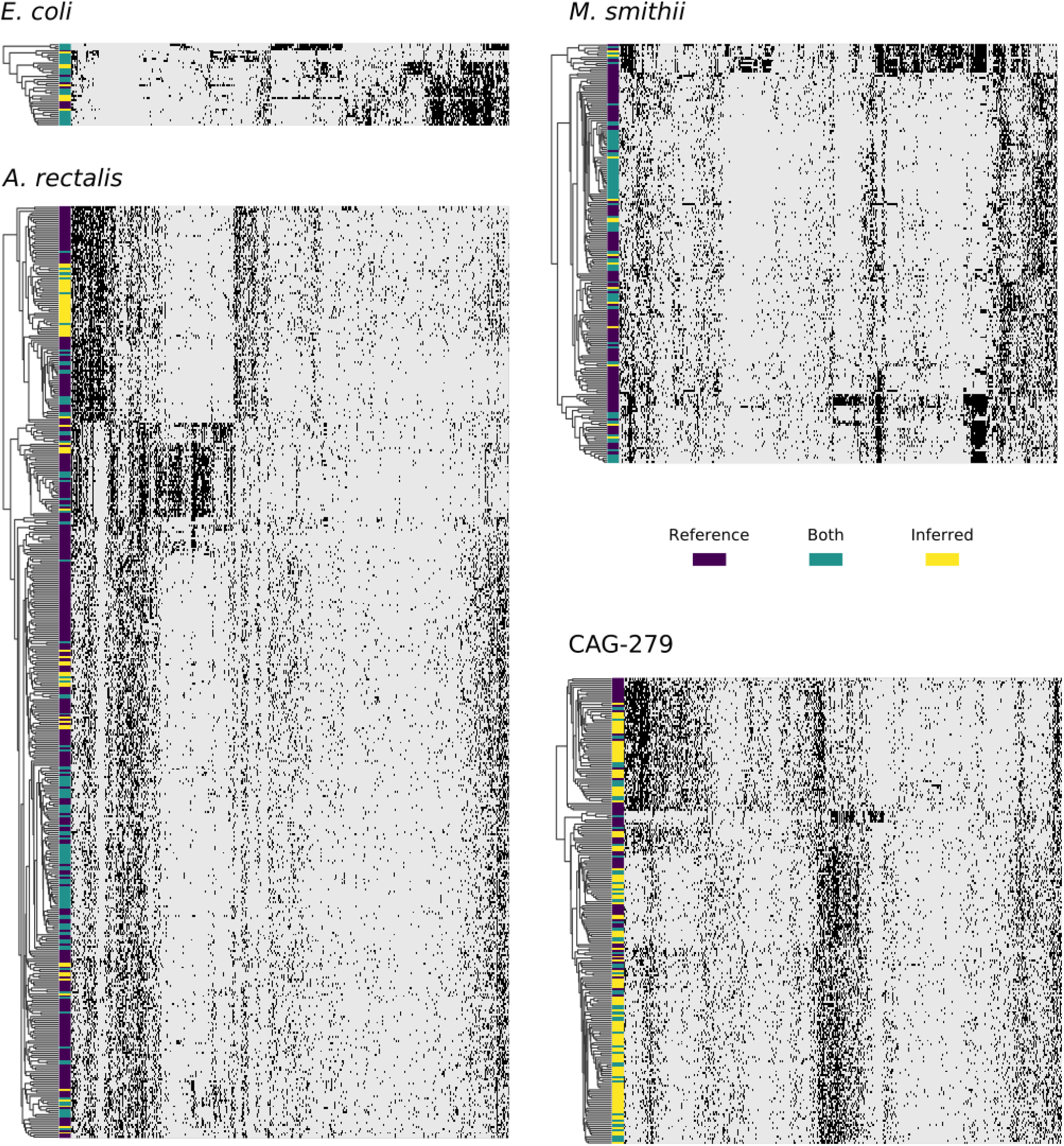
Concordance between reference and StrainFacts inferred strain genotypes for four prevalent species in the human gut microbiome. Heatmap rows represent consensus genotypes from co-clustering of reference and inferred strains and columns are 3500 randomly sampled SNP sites (grey: reference and black: alternative allele). Colors to the left of the heatmap indicate clusters with only reference strains (dark purple), only inferred strains (yellow), or both (teal). Rows are ordered by hierarchical clustering built on distances between consensus genotypes and columns are ordered arbitrarily to highlight correlations between SNPs.

For *E. coli*, we identified 40 strain clusters with 93% of inferred strains and 94% of references falling into clusters containing strains from both sources (“shared” clusters), which is significantly more overlap than expected after random shuffling of cluster labels (p=0.002 by permutation test). While most metagenome-inferred genotypes are similar to those found in genome reference databases, we observed some clusters composed only of StrainFacts strains, representing novel lineages. However, these strains are no more common than after random permutation (p=0.81), matching our expectations for this well-studied species.

We next asked if these trends hold for the other species. While *A. rectalis* had a much greater number of clusters (456), 69% of inferred strains and 45% of reference strains are nonetheless found to be in shared clusters, significantly more than would be expected with random shuffling of cluster labels (p=0.002 by permutation test). Correspondingly, we do not find evidence for enrichment of inferred strains in novel clusters (p=0.71). We find similar results for *M. smithii* and CAG-279—the fraction of strains in shared clusters is significantly greater than after random reassignment (p<0.001 for both), and there is no evidence for enrichment of inferred strains in novel clusters (p = 1.0 for both). Overall, the concordance between reference and inferred strains supports not only the credibility of StrainFacts’ estimates, but also suggests that our *de novo* inferences capture a substantial fraction of previously documented strain diversity, even in well studied species.

Going beyond the extensive overlap of strains with reference genomes and StrainFacts inferences, we examined clusters in which references are absent or relatively rare. Visualizing a dendrogram of consensus genotypes from co-clustered strains (Fig. 4) we observe some sections of the *A. rectalis* dendrogram with many novel strains. Similarly, for CAG-279, the sheer number of inferred strains relative to genomes in reference databases means that fully half of all genotype clusters are entirely novel, emphasizing the power of StrainFacts inferences in understudied species. Future work will be needed to determine if these represent new subspecies currently missing from reference databases.

### Species inhabiting the human gut exhibit distinct biogeography observed across independent metagenomic studies

Large metagenomic collections allow us to examine intraspecific microbial diversity at a global scale and among dozens of studies. Towards this end, we identified the most abundant StrainFacts strain of *E. coli, A. rectalis, M. smithii*, and CAG-279 in stool samples across 34 independent studies. Across all four species, we observe some strains that are distributed globally as well as others that appear specific to one country of origin (Fig. 5, Supplementary Fig. S5). For example, among the 198 dereplicated, inferred strains of *A. rectalis*, 75 were found as the dominant strain (i.e. highest relative abundance) in samples collected on three or more continents. While this makes it challenging to consistently discern where a sample was collected based on its dominant strain of a given species, we nonetheless find that studies with samples collected in the United States of America form a distinct cluster, as do those from China, and the two are easily distinguished from one another and from most other studies conducted across Europe and North America (Fig. 5). Our observation of a distinct group of *A. rectalis* strains enriched in samples from China is consistent with previous results (Scholz et al., 2016; Costea et al., 2017a; Truong et al., 2017).

**Figure 5:**
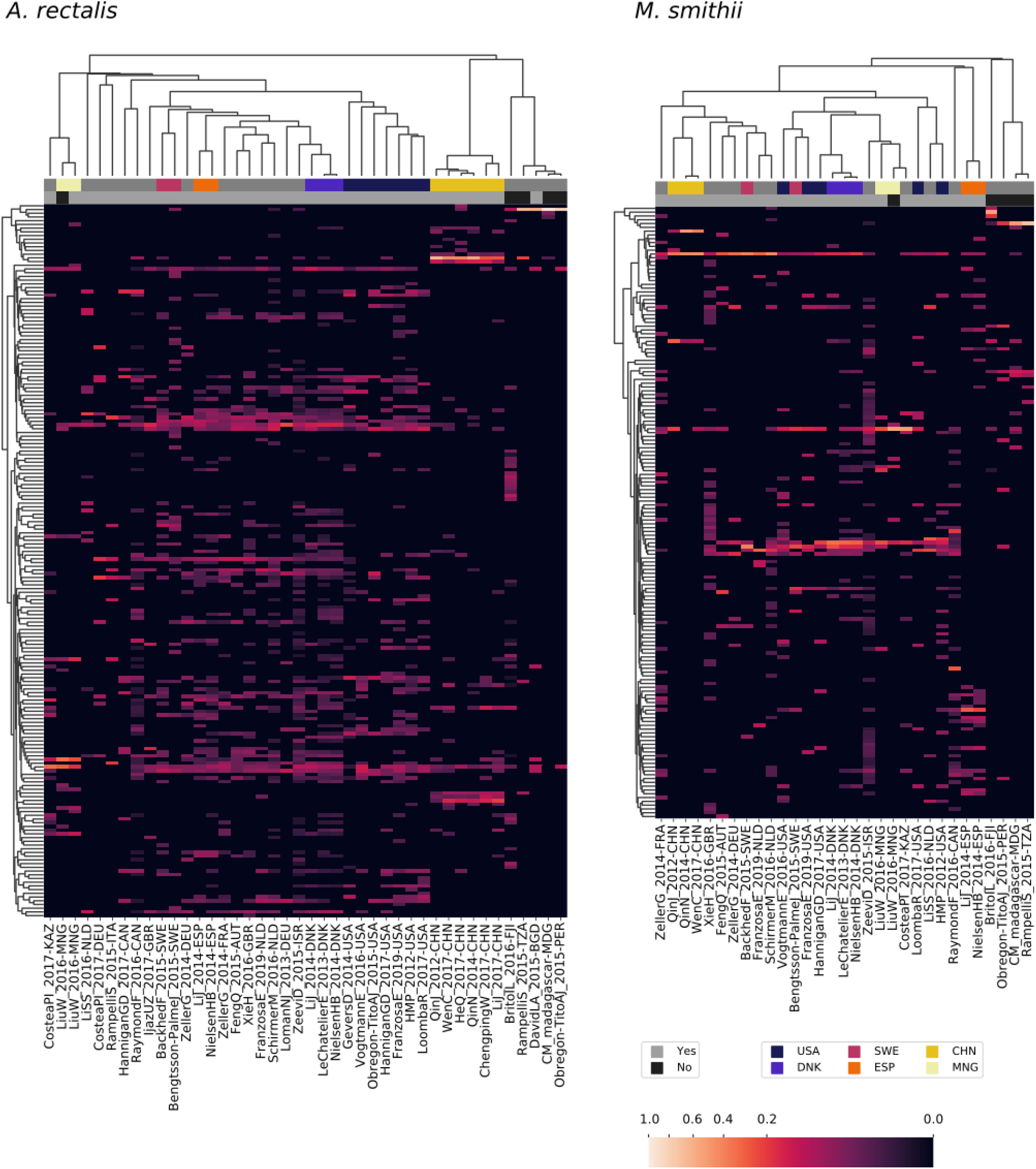
Patterns in strain dominance according to geography and lifestyle across thousands of publicly available metagenomes in dozens of independent studies for two common members of the human gut microbiome. Columns represent collections of samples from individual studies and are further segmented by country and lifestyle (westernized or not). Rows represent strains inferred by StrainFacts. Cell colors reflect the fraction of samples in that study segment with that strain as the most abundant member. Study segments are omitted if they include fewer than 10 samples. Row ordering and the associated dendrogram reflect strain genotype distances, while the dendrogram for columns is based on their cosine similarity. Studies with samples collected in several countries with notable clustering for one or more species are highlighted with colors above the heatmap. Additionally, studies from westernized populations are indicated. Both a study identifier and the ISO 3166-ISO country-code are included in the column labels.

These general trends hold across the other three species. In *M. smithii*, independent studies in the same country often share very similar strain dominance patterns (e.g. see clustering of studies performed in each of China, Mongolia, Denmark, and Spain in Fig. 5). In *E. coli*, while many strains appear to be distributed globally, independent studies from China still cluster together based on patterns in strain dominance (see Supplementary Fig. S5). Notably, in CAG-279, studies with individuals in westernized societies do not cluster separately from the five other studies, suggesting that host lifestyle is not highly correlated with specific strains of this species. The variety of dominance patterns across the four species described here suggest that different mechanisms may drive intraspecific biogeography depending on the biology and natural history of the species. As the coverage of diverse microbiomes grows, StrainFacts will enable future studies disentangling the contributions of lifestyle, dispersal limitation and other factors in the global distribution of strains.

### Linkage disequilibrium decay suggests variation in recombination rates across microbial species

Studying the population genetics of host-associated microbes has the potential to elucidate processes of transmission, diversification, and selection with implications for human health and perhaps even our understanding of human origins (Linz et al., 2007; Garud and Pollard, 2019). To demonstrate the application of StrainFacts to the study of microbial evolution, we examined patterns in pairwise LD, here calculated as the squared Pearson correlation coefficient (r^2^). This statistic can inform understanding of recombination rates in microbial populations (Vos, 2009; Garud et al., 2019). Genome-wide, LD, summarized as the 90th percentile r^2^ (LD_90_, Vos et al., 2017), was substantially higher for *E. coli* (0.24) than *A. rectalis* (0.04), *M. smithii* (0.11), or CAG-279 (0.04), perhaps suggesting greater population structure in the species and less panmictic recombination.

We estimated LD distance-decay curves for SNPs, using a single, high-quality reference genome for each species to obtain estimates of pairwise distance between SNP sites. For all four species, adjacent SNPs were much more tightly linked, with an LD_90_ of >0.999. LD was still dramatically above background at 50 bases of separation, and fell rapidly with increasing distance (Fig. 6). The speed of this decay was different between species, which we characterized with the LD_½,90_: the distance at which the LD_90_ was less than 50% of the value for adjacent SNPs (Vos et al., 2017). *M. smithii* exhibited by far the slowest decay, with a LD_½,90_ of 520 bases, followed by *E. coli* at 93 bases, CAG-279 at 66, and *A. rectalis* at just 28 bases. This variation in decay profiles may reflect major differences in recombination rates across populations.

**Figure 6:**
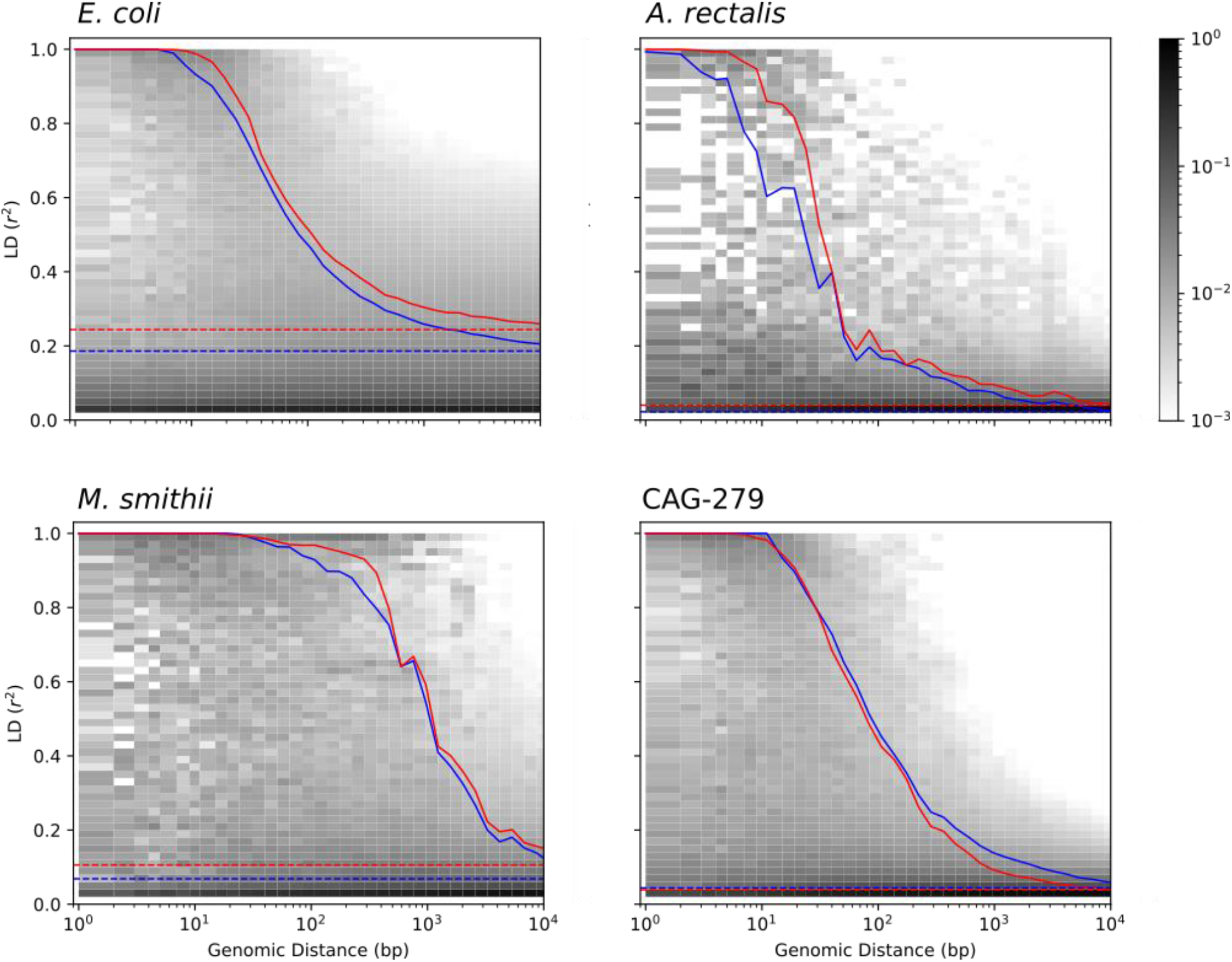
Pairwise LD across genomic distance estimated from inferred genotypes for four species. LD was calculated as r^2^ and genomic distance between polymorphic loci is based on distances in a single, representative genome. The distribution of SNP pairs in each distance window is shown as a histogram with darker colors reflecting a larger fraction of the pairs in that LD bin, and the LD_90_ for pairs at each distance is shown for inferred strains (red), along with an identical analysis on strains in the reference database (blue). Genome-wide LD_90_ (dashed lines) is also indicated.

To validate our findings, we ran identical analyses with dereplicated genotypes from genomes in the GT-Pro reference database. Estimates of both LD and the distance-decay relationship are very similar between inferred and reference strains, reinforcing the value of genotypes inferred from metagenomes for microbial population genetics. Interestingly, for three of the four species (*E. coli, A. rectalis*, and *M. smithii*), LD estimates from StrainFacts strains were significantly higher than from references (p<1e-10 for all three by Wilcoxon test), while CAG-279 exhibited a trend towards the reverse (p=0.85). It is not clear what might cause these quantitative discrepancies, but they could reflect differences in the set of strains in each dataset. Future studies expanding this analysis to additional species will identify patterns in recombination rates across broader microbial diversity.

## Discussion

Here we have described StrainFacts, a novel tool for strain inference in metagenomic data. StrainFacts models metagenotype data using a fuzzy-genotype approximation, allowing us to estimate both the relative abundance of strains across samples as well as their genotypes. To accelerate analysis compared to the current state-of-the-art, we harness the differentiability of our model to apply modern, gradient-based optimization and GPU-parallelization. Consequently, StrainFacts can scale to tens-of-thousands of samples while inferring genotypes for hundreds of strains. On simulated benchmarks, we show that StrainFacts has comparable accuracy to Strain Finder, and we validate strain inferences *in vivo* against genotypes observed by single-cell genomics. Finally, we apply StrainFacts to a database of tens of thousands of metagenomes from the human microbiome to estimate strains *de novo*, allowing us to characterize strain diversity, biogeography, and population genetics, without the need for cultured isolates.

Beyond Strain Finder, other alternatives exist for strain inference in metagenomic data. While we do not directly compare to DESMAN, runtimes of several hours have been reported for that tool on substantially smaller simulated datasets (Quince et al., 2017), and hence we believe that StrainFacts is likely the most scalable implementation of the metagenotype deconvolution approach. Still other methods apply regularized regression (e.g. Lasso Albanese and Donati, 2017) to decompose metagenotypes—essentially solving the abundance half of the deconvolution problem but not the genotypes half—or look for previously determined strain signatures (e.g. k-mers Panyukov et al., 2020) based on known strain genotypes. However, both of these require an *a priori* database of the genotypes that might be present in a sample. An expanding database of strain references can make these approaches increasingly useful, and StrainFacts can help to build this reference.

Several studies avoid deconvolution by directly examining allele frequencies inferred from metagenotypes. While consensus (Truong et al., 2017; Zolfo et al., 2017) or phasing (Garud et al., 2019) approaches can accurately recover genotypes in some cases, their use is limited to low complexity datasets, with sufficient sequencing depth and low strain heterogeneity. Likewise, measuring the dissimilarity of metagenotypes among pairwise samples indicates shared strains (Podlesny and Fricke, 2020), but this approach risks confounding strain mixing with genotype similarity. Finally, assembly (Li et al., 2019) and read-based methods (Cleary et al., 2015) for disentangling strains are most applicable when multiple SNPs can be found in each sequencing read. With ongoing advancements in long-read (Vicedomini et al., 2021) and read-cloud sequencing (Kuleshov et al., 2016; Kang et al., 2018), these approaches will become increasingly feasible. Thus, StrainFacts occupies the same analysis niche as Strain Finder and DESMAN, and it expands upon these reliable approaches by providing a scalable model fitting procedure.

Fuzzy genotypes enable more computationally efficient inference by eliminating the need for discrete optimization. Specifically, we used well-tested and optimized gradient descent algorithms implemented in PyTorch for parameter estimation. In addition, fuzzy genotypes may be more robust to deviations from model assumptions. For instance, an intermediate genotype could be a satisfactory approximation when gene duplications or deletions are present in some strains. While we do not explore the possibility here, fuzzy genotypes may provide a heuristic for capturing uncertainty in strain genotypes. Future work could consider propagating intermediate genotype values instead of discretizing them.

StrainFacts builds on recent advances in metagenotyping, in particular our analyses harnessed GT-Pro (Shi et al., 2021) to greatly accelerate SNP counting in metagenomic reads. While we leave a comparison of StrainFacts performance on the outputs of other metagenotypers to future work, StrainFacts itself is agnostic to the source of input data. It would be straightforward to extend StrainFacts to operate on loci with more than two alleles or to use metagenotypes from a tool other than GT-Pro. It would also be interesting to extend StrainFacts to use SNPs outside the core genome that vary in their presence across strains.

Combined with the explosive growth in publicly available metagenomic data and the development of rapid metagenotyping tools, StrainFacts enables the *de novo* exploration of intraspecific microbial diversity at a global scale and on well-powered cohorts with thousands of samples. Future applications could examine intraspecific associations with disease, track the history of recombination between microbial subpopulations, and measure the global transmission of host-associated microbes across human populations.

## Acknowledgements

Barbara Engelhardt provided valuable feedback on this project.

## Author Contributions

- BJS: conceptualization, data curation, formal analysis, methodology, software, visualization, writing - original draft, writing - review & editing
- XL: investigation, data curation, methodology, writing - original draft, writing - review & editing
- ZJS: data curation, methodology, writing - review & editing
- AA: funding acquisition, supervision
- KSP: conceptualization, writing - review & editing, funding acquisition, supervision

## Data Availability Statement

Metagenomic and single-cell sequencing data from the FMT study will be uploaded to the SRA under BioProject PRJNA737472. Publicly available metagenomes are available under various other accessions described in (Shi et al., 2021). Strain genotypes from the GT-Pro reference database are publicly available at https://fileshare.czbiohub.org/s/waXQzQ9PRZPwTdk. All other code and metadata needed to reproduce these results are available at https://github.com/bsmith89/StrainFacts-manuscript.

## Supplementary Materials

### Supplementary Methods

**Figure S1:**
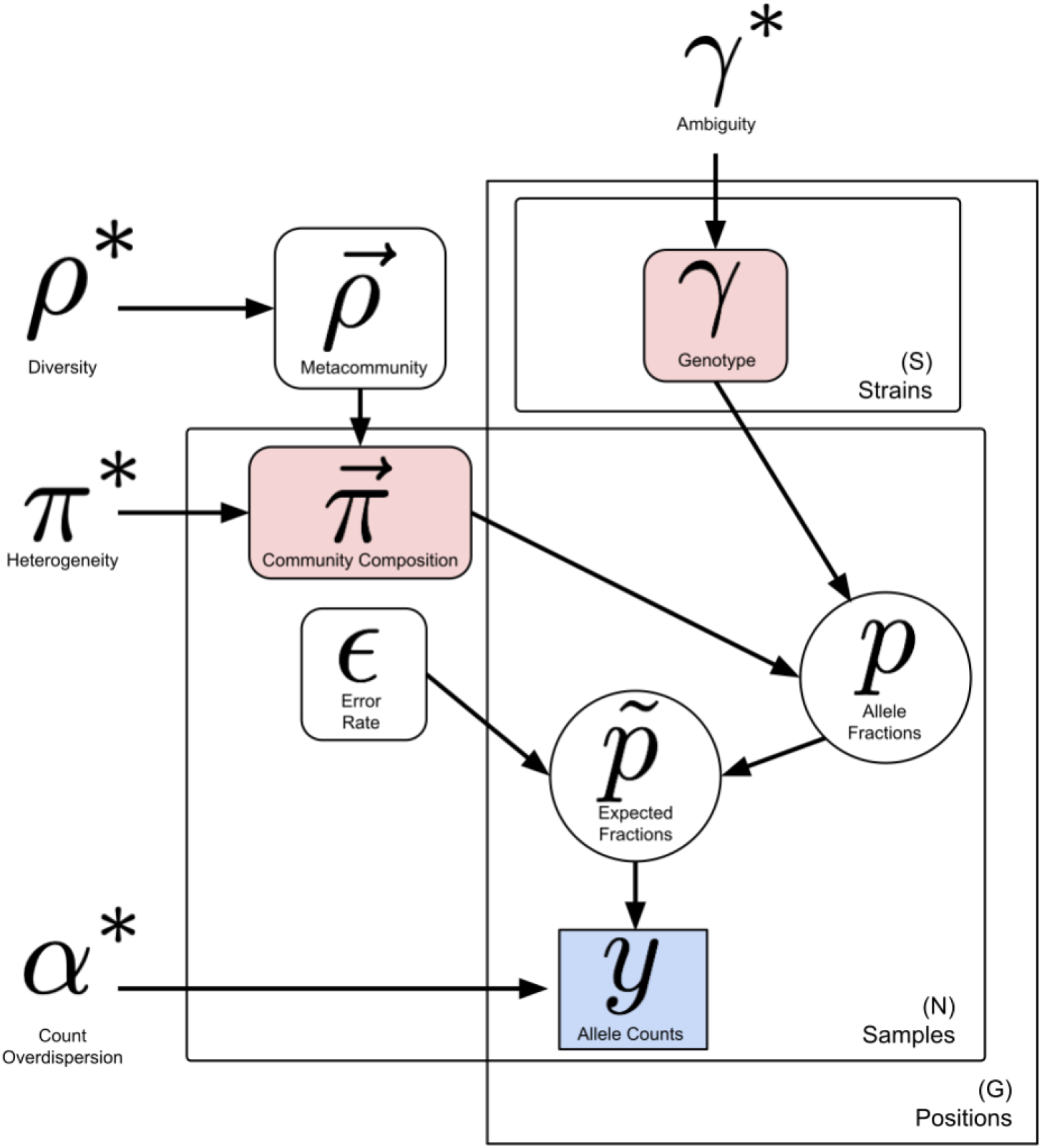
Graphical representation of the StrainFacts model including hyperparameters. Symbols include observed data (blue box), deterministic terms (circles), key parameters being estimated (red boxes), and key hyperparameters (unenclosed). Plates behind terms indicate the dimensionality and indexing of the variables and arrows connect the terms that directly depend on one another.

#### The shifted, scaled Dirichlet distribution

The k-dimensional SSD is a 2k + 1 parameter family which includes the Dirichlet distribution as a special case, and is defined by Aitchenson “perturbation” (⊕) and “powering” (⊙) operations (Aitchison, 1986) applied to a Dirichlet-distributed random variable. Given a random variable **X** ∼ Dirichlet(α), α ∈ ℝ^*K*^, **X** ∈ 𝒮^*D*^ if **Y = p** ⊕ (*a* ⊙ **X**), *a* ∈ ℝ_+_, **p** ∈ 𝒮^*K*^, **Y** ∈ 𝒮^*K*^ then **Y** ∼ SSD(**α, p**, *a*).

In this work, we limit our use of this distribution to **α = 1**, i.e. **X** distributed uniformly on the K-simplex before powering and perturbation. For values of *a* > 1, the probability mass shifts towards the edges of the simplex, and we use this property in order to induce sparsity in our estimates.

#### Parameter initialization and optimization

In select initial values of **Π** and **Γ** using an NMF based approach. First, we transform metagenotypes from counts to an *N* × *G* × 2 matrix of allele frequencies, and stack this into an *N* × 2*G* matrix, **P**′, with columns of reference alleles followed by columns of alternative alleles. We then use canonical NMF—implemented in the scikit-learn library (Pedregosa et al., 2011)— to factorize this data matrix into **Π**′ and **Γ**′, where **P**′ ≈ **Π**′ × **Γ**′ with a shared, inner dimension of size *S*. After reversing the stacking of alleles, we get back a matrix 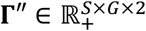. Since **Π**′ and **Γ**″ likely do not conform to the constraints of strain deconvolution, we transform them into initial values as follows: 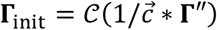 and 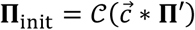 where 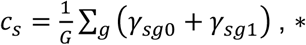 denotes element-wise multiplication, and 𝒞(⋅) is normalization over the last dimension to the standard simplex (i.e. summing-to-one).

Model parameters are transformed to the unconstrained space using Pyro’s built-in defaults. Parameters other than **Π** and **Γ** are initialized randomly to a point on the interval (−2,2) in the transformed space. We then apply the Adamax algorithm for stochastic gradient descent using an initial learning rate `lr_init`. To increase the probability of finding a global maximum, we take a prior annealing approach (Neuwald and Liu, 2004): for an initial `n_wait` number of steps of the optimization routine, the hyperparameters *γ*^*^ and *ρ*^*^ are set to initial values with less stringent regularization and are then exponentially relaxed to their final values during the next `n_anneal` steps. After this annealing period, we continue taking gradient steps until the value of the loss function has not improved for 100 steps, at which point we halve the learning rate. Optimization is stopped when the learning rate falls below a minimum threshold, `lr_min`.

#### Fitting full length genotypes

Because many metagenotypes had more than 5,000 SNP sites, we use a refitting approach to get full length strain genotypes. This is accomplished by conditioning our model on both the observed data and the previously estimated **Π**. In addition, we update the value of two of the hyperparameters; we set *γ*^*^ **=** 1.0, and *α*^*^ **=** 200. After refitting the other parameters, this results in a new estimate of **Γ**. Since SNPs are statistically separable when **Π** is conditioned out, this allows us to iteratively refit arbitrary subsets of SNPs before recombining them into a full length genotype matrix.

#### Single-cell genomic sequencing

##### Cell isolation from stool samples

Bacterial cells were isolated from fecal samples according to previously published protocol (Hevia et al., 2015) with modifications. Briefly, 0.2-0.5 g of fecal samples were homogenized in 10 mL of PBS buffer by vertexing. After filtering with a 50 μm cell strainer (Corning, 431752) to remove most of the fecal particles, the flow through suspension was loaded on top of 3.5 mL of 80% Nycodenz® (Cosmo Bio USA, AXS-1002424) in a 15 mL conical tube. The tube was centrifuged for 40 min at 4 °C (4700 x g). The layer corresponding to microbiota was extracted and washed with PBS for 3 times. The cells were directly processed for hydrogel encapsulation or stored in DNA/RNA shield (Zymo Research, R1100-50) at -80 °C for long term storage.

##### High-throughput single bacterial sequencing

Barcoded single cell bacterial sequencing libraries were constructed by modified SiC-protocol Seq leveraging Mission Bio Tapestri (Lan et al., 2017).

##### Cell encapsulation in hydrogel beads

Cell suspension (100 million per mL in PBS, 500 μL) was mixed with 500 μL of polyacrylamide precursor solution with 12% acrylamide(Thermo Scientific, AAJ62480AP), 1% N,N′-bis(acryloyl)cystamine (Sigma, A5912), 20 mM Tris (pH 8.0), 0.6% sodium persulfate (Sigma, 216232), and 20 mM NaCl. After adding 1mL of HFE 7500 with 2% surfactant (008-FluoroSurfactant, RanBiotechnologies), heterogenous droplets were generated by passing the oil/aqueous mixture through a syringe with 23.5 G blunt needle for 5 time. 20 μL of N,N,N′,N′-tetramethylethylenediamine (Sigma, 411019) was added into the emulsion and the emulsion was incubated at 70 °C for 30 min and at room temperature for 1 hour for gelation. The emulsion was centrifuged at 1000xG for 10 min and the oil layer was removed by a gel loading tip. To the hydrogel layer, 1 mL of 20% PFO (1H,1H,2H,2H-perfluoro-1-octanol, Sigma, 370533) in HFE 7500 and the mixture was vortexed and shaking for 10 min to break the emulsions. After centrifugation at 1000xG for 10 min, PFO were removed, and the hydrogel beads were then washed with PBS with 0.4% tween 20 for 3 times. The beads were then resuspended in 40% sucrose in PBS with 0.4% tween 20. A differential velocity centrifugation was performed to select the hydrogel beads within the size between 5 to 25 μm.

##### Cell lysis and DNA purification in hydrogel beads

100 μL of beads were treated in a solution of 1 mL TE buffer solution containing 2.5 mM EDTA (Teknova), 10mM NaCl (Sigma-Aldrich), 2U zymolyase, 5 U Lysostaphin, 50 U mutanolysin, and 20 mg Lysozyme at 37 °C overnight. The lysate mixture was then centrifuged at 3000 xG for 3 min, the supernatant removed, and 1 mL of TE buffer with 4U of Proteinase K, 1% triton X100 and 100 mM of NaCl was added to digest cellular proteins. The solution is incubated at 50 °C for 30 min. Following lysis, the beads was thoroughly washed to ensure complete removal of detergents and other chemicals which may inhibit the downstream reactions. The washes were performed in 10 mL volume with centrifugation magnitudes of 3000 x g between washes.

##### Tagmentsation reagents

25 μL Blocked ME Complement /5Phos/C*T* G*T*C* T*C*T* T*A*T* A*C*A*/3ddC/ (200 nM, IDT), 25 μL Tn5-Fwd-oligo GTACTCGCAGTAGTCAGATGTGTATAAGAGACAG (100 nM, IDT), and 25 μL Tn5-Rev-oligo TACCCTTCCAATTTAACCCTCCAAGATGTGTATAAGAGACAG (100 nM, IDT) and 25 μLTris buffer were mixed well in a PCR tube by pipetting. The mixture was incubated on a thermal cycler using the following program: 85°C for 2 min, cools to 20 °C with a ramping rate at 0.1 °C/s, 20 °C for 1 min, then hold at 4 °C with lid at 105°C. 100 μL of glycerol was then added into the annealed oligo. The unloaded Tn5 (1 mg/mL, expressed by QB3 MacroLab, Berkeley, CA.) was diluted at a 1:1 ratio in the Illumina dilution buffer (50% Glycerol, 100 mM NaCl, 0.1 mM EDTA, 1 mM DTT, and 0.1% NP40 in 50 mM Tris-HCl pH 7.5 buffer), followed by mixing at a 1:1 ratio with the pre-annealed adapter/glycerol mix. The mixture was incubated at room temperature for 30 min then stored at -20 °C.

##### Single cell DNA tagmentaion

The beads were resuspended in the density matching buffer (10 mM MgCl2, 1% NP40, and 17% Optiprep in 20 mM TAPS pH 7.0 buffer) to the final cell density of 3000 cells/μL. 25 μL of assembled Tn5 was mixed with 25 μL of tagmention buffer (10 mM MgCl2, 10 mM DTT in 20 mM TAPS pH 7.0 buffer). The MissionBio Tapestri DNA cartridge and a 0.2 mL PCR tube were mounted onto the Tapestri instrument. 50 μL of the beads in density matching buffer was loaded into the reservoir 2 (the reservoir for cell suspension), 50 μL of the Tn5 in tagmentation buffer was loaded into reservoir 1 (the reservoir for lysis buffer), and 200 μL of Encapsulation oil was loaded into reservoir 3. After applying the DNA gasket on top of the cartridge and closing the instrument lid, droplets were generated by running the Step1: Encapsulation program. The droplets were incubated at 37°C for 60 min and then 50°C for 30 min.

##### Barcoding PCR

Barcoding droplet PCR were performed according to the MissionBio Tapestri protocol with minor modification. 8 PCR tubes and DNA cartridge were mounted onto the Tapestri instrument. 200 μL and 500 μL of Electrode solution were loaded into reservoirs 4 and 5 respectively. After running the Priming program, 5 μL of reverse barcoding oligo was mixed with 295 μL MissionBio Barcoding Mix and loaded into reservoir 8 of the DNA cartridge. The droplets from previous step (∼80 uL), 200 μL of barcoding beads, and 1.25 mL of Barcoding oil were loaded into reservoir 6, 7 and 9, respectively. After applying the DNA gasket on top of the cartridge and closing the lid, the droplets were merged with barcoding beads and PCR reagents by running the Cell Barcoding program on the Tapestri instrument. The droplets collected in the 8 PCR tubes were treated with UV for 8 min (Analytik Jena Blak-Ray XX-15L UV light source) and the bottom layer of oil in each tube were removed using a gel loading tip to leave up to 100 μL of droplets. The tubes were then thermo-cycled on a PCR instrument with the following program: 10 min at 72°C for 1 cycle, 3 min at 95°C for 1 cycle, (15 s at 95°C, 15 s for 55°C, and 2 min at 72°C) for 20 cycles, and 5 min at 72°C for 1 cycle.

##### Sequencing library preparation

The thermal cycled droplets in the 8 PCR tubes were carefully transferred into two 1.5 mL centrifuge tubes (4 PCR tube content in each). If there were merged droplets present, they were carefully removed using a 2 μL pipette. 20 μL PFO were added into each tube and mixed well by vortex. After centrifuging the top aqueous layers in each tube was transferred into new 1.5 mL tubes and water was added to bring the total volume to 400 μL. The barcoding product was purified using 0.7X Ampure XP beads (Beckman Coulter, A63882) and eluted into 60 μL H2O and stored at -20°C until next step. The concentrations of the barcoding product were measured with Qubit™ 1X dsDNA Assay Kits (ThermoFisher, Q33230).

The sequencing library were then prepared by attaching P5 and P7 sequences to the barcoding products using Nextera primers. The library PCR reactions were performed with 25 uL Kapa HiFi Master mix 2X, 5 uL Library P5 index primer (4 uM), 5 uL Library P7 index primer (4 uM), 10 uL purified barcoding products (normalized to 0.2 ng/uL), and 5 uL of nuclease free water. The PCR tubes were thermal cycled with the following program: 3 min at 95°C for 1 cycle, (20 s at 98°C, 20 s for 62°C, and 45 s at 72°C) for 10 cycles, and 2 min at 72°C for 1 cycle. The sequencing library was purified with 0.69X Ampure XP beads and eluted into 12 uL nuclease-free water. The library was quantified with Qubit™ 1X dsDNA Assay Kits and DNA HS chips on bioanalyzer or D5000 ScreenTape (Agilent, 5067-5588) on Tapestation (Agilent, G2964AA). The libraries were pooled and paired-end sequenced by Novogene with a partial lane on Illumina NovaSeq 6000.

##### Single cell read files preparation

Sequencing data were processed using a custom python script (`mb_barcode_and_trim.py`) available on GitHub at https://github.com/AbateLab/MissonBioTools. For all reads, combinatorial cell barcodes were parsed from Read 1, using cutadapt (v2.4) and matched to a barcode whitelist. Barcode sequences within a Hamming distance of 1 from a whitelist barcode were corrected. Reads with valid barcodes were trimmed with cutadapt to remove 5′ and 3′ adapter sequences and demultiplexed into single-cell FASTQ files by barcode sequences using the script `demuxbyname.sh` from the BBMap package (v.38.57).

### Supplementary Results

**Figure S2:**
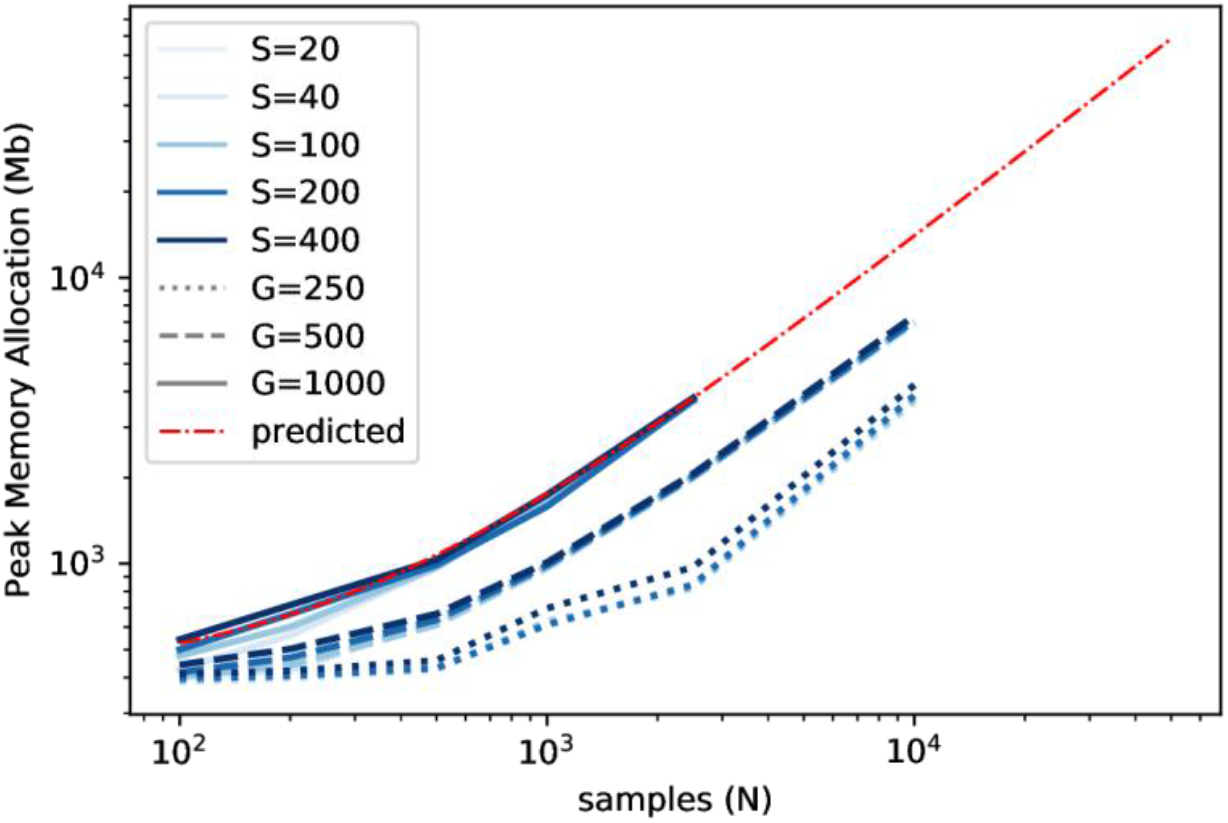
Maximum memory allocation across varying numbers of strains (S, line shade), SNPs (G, line style), and samples is plotted for StrainFacts models. Median of 9 replicate runs is shown. Maximum memory requirements are extrapolated to higher numbers of samples for a model with 1000 SNP sites (red line). An abridged version of this plot is included as Fig. 1.

**Figure S3:**
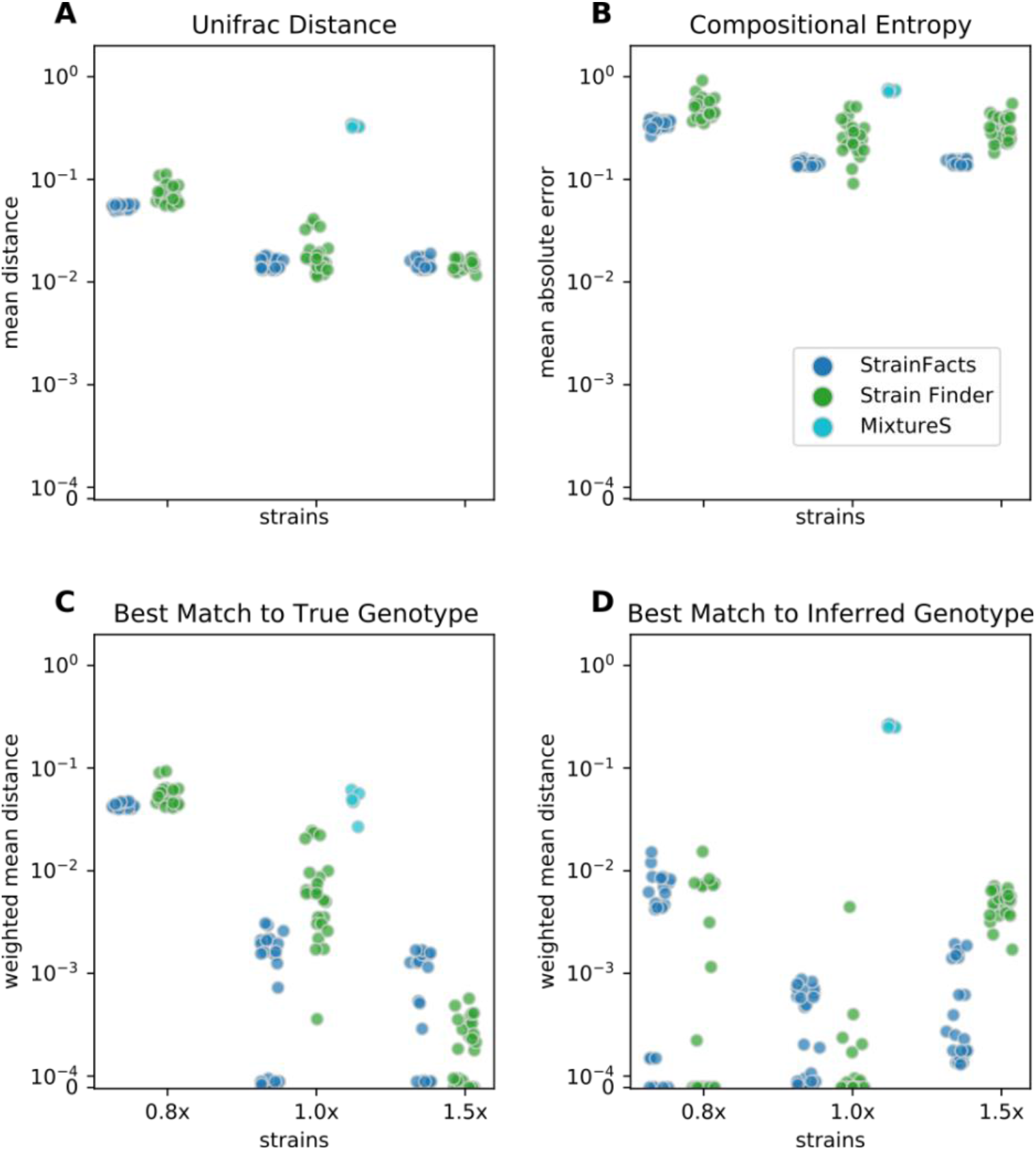
Extension of accuracy evaluation for StrainFacts and Strain Finder with additional results for MixtureS. Results are identical to panels A, C, D, and E in Fig. 2 (here panels **A**-**D**, respectively). Simulations are shown for five simulations with 250 SNP positions, 200 samples, and 40 strains. While StrainFacts and Strain Finder each have 32, 40, and 60 strains specified (the 0.8x, 1.0x, 1.5x parameterizations), MixtureS does not specify the number of strains a priori, and points are arbitrarily placed with the 1x parameterization. Similarly, MixtureS runs are deterministic; hence only one fit for each of the five simulations is shown.

**Figure S4:**
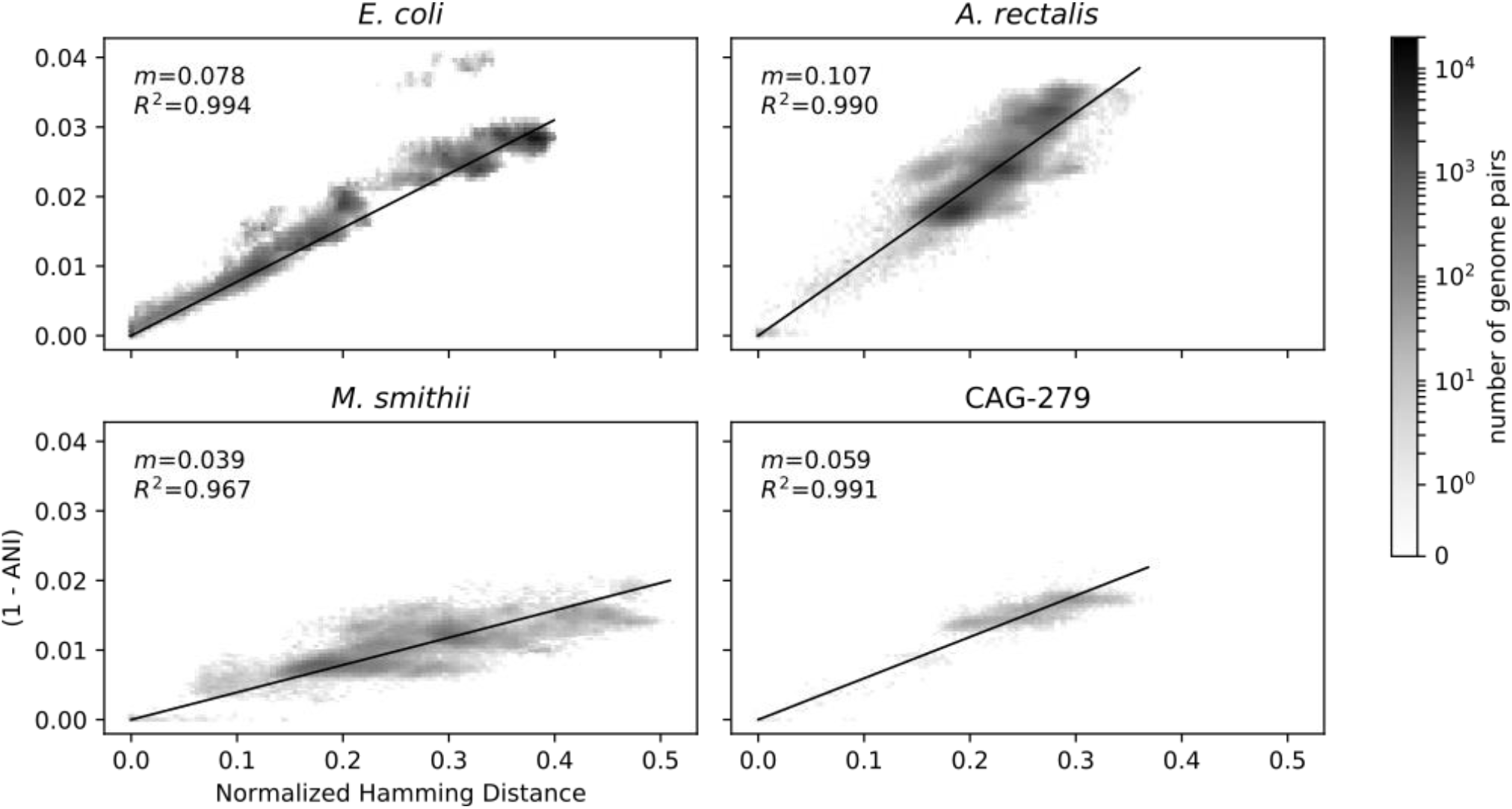
Emipirical relationship between ANI and genotype distance among reference genomes in the GT-Pro database. Genotype distance is defined as the normalized Hamming distance at SNP sites considered by GT-Pro. All pairwise genome comparisons are plotted as a 2D histogram, with greater density indicated with darker colors. For each species, a linear regression calculated without an intercept term is shown (black line), and the constant of proportionality and uncentered R^2^ is also indicated.

**Figure S5:**
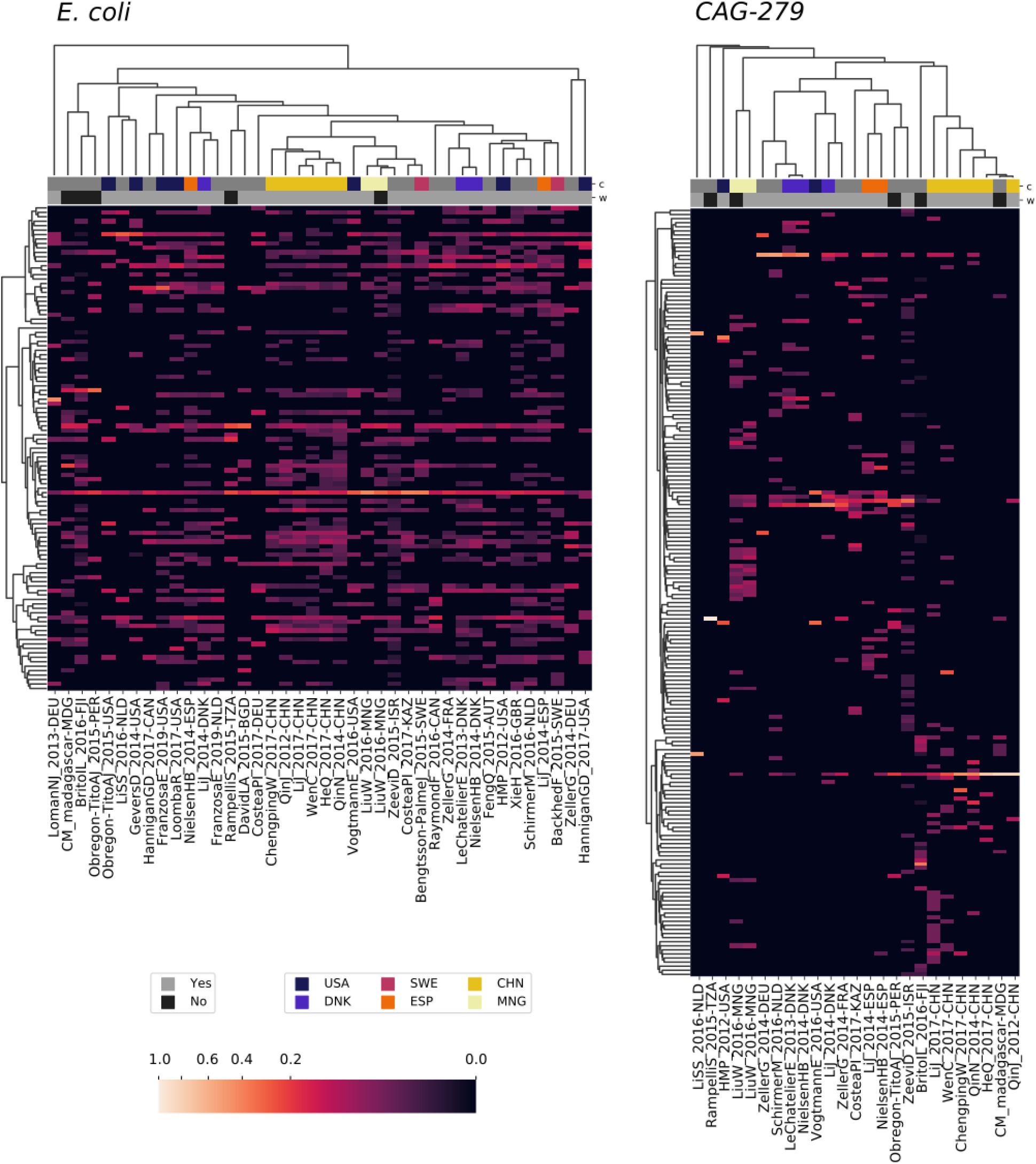
Patterns in strain dominance according to geography and lifestyle across thousands of publicly available metagenomes in dozens of independent studies for two additional members of the human gut microbiome. Visual elements are identical to Fig. 5: Columns represent collections of samples from individual studies and are further segmented by country and lifestyle (westernized or not). Rows represent strains inferred by StrainFacts. Cell colors reflect the fraction of samples in that study segment with that strain as the most abundant member. Study segments are omitted if they include fewer than 10 samples. Row ordering and the associated dendrogram reflect strain genotype distances, while the dendrogram for columns is based on their cosine similarity. Colors above the heatmap reflect the country in which samples were collected as well as whether samples were collected from individuals with a westernized lifestyle. Both a study identifier and the ISO 3166-ISO country-code are included in the column labels.

